# A pharmacoproteomic landscape of organotypic intervention responses in Gram-negative sepsis

**DOI:** 10.1101/2022.08.29.503941

**Authors:** Tirthankar Mohanty, Christofer A. Q. Karlsson, Yashuan Chao, Erik Malmström, Eleni Bratanis, Andrietta Grentzmann, Martina Mørch, Victor Nizet, Lars Malmström, Adam Linder, Oonagh Shannon, Johan Malmström

**Author notes:** Correspondence: Johan Malmstrom, Oonagh Shannon. These authors contributed equally to the work (Tirthankar Mohanty, Christofer A. Q. Karlsson).

## Abstract

Sepsis is the major cause of mortality across intensive care units globally, yet details of accompanying pathological molecular events remains unclear. This knowledge gap has resulted in ineffective development of sepsis-specific biomarkers and therapies, and suboptimal treatment regimens to prevent or reverse organ damage. Here, we used pharmacoproteomics to score treatment effects in a murine *Escherichia coli* sepsis model based on changes in the organ, cell, and plasma proteome landscapes. A combination of pathophysiological read-outs and time-resolved proteome maps of organs and blood enabled us to define time-dependent and organotypic proteotypes of dysfunction and damage upon administration of several combinations of the broad-spectrum beta-lactam antibiotic meropenem (Mem) and/or the immunomodulatory glucocorticoid methylprednisolone (Gcc). Three distinct response patterns were identified, defined as intervention-specific reversions, non-reversions, and specific intervention-induced effects, which depended on the underlying proteotype and varied significantly across organs. In the later stages of the disease, Gcc enhanced some positive treatment effects of Mem with superior reduction of the inflammatory response in the kidneys and partial restoration of sepsis-induced metabolic dysfunction. Unexpectedly, Mem introduced sepsis-independent perturbations in the mitochondrial proteome that were to some degree counteracted by Gcc. In summary, this study provides a pharmacoproteomic resource describing the time-resolved septic organ failure landscape across organs and blood, coupled to a novel scoring strategy that captures unintended secondary drug effects as an important criterion to consider when assessing therapeutic efficacy. Such information is critical for quantitative, objective, and organotypic assessment of benefits and unintended effects of candidate treatments in relationship to dosing, timing, and potential synergistic combinations in murine sepsis models.

## Introduction

Sepsis is a leading cause of mortality in intensive care units (Reinhart et al., 2017; Sakr et al., 2018), and plays a contributing role in up to 19% of deaths worldwide (Rudd et al., 2020). Recent Sepsis-3 guidelines define the syndrome as life-threatening organ dysfunction caused by a dysregulated host response to infection (Singer et al., 2016). Sepsis is remarkably variable in its manifestations, making it challenging to diagnose, predict disease progression, and treat. A wide range of pathogens are implicated, each with unique inherent abilities to evade immune responses and cause tissue injury. Equally, innate and adaptive immune response genes are among the most polymorphic in the human genome, such that individual or groups of host genotypes respond differently to the same infectious challenge.

Sepsis is characterized by dynamic disease changes, progression from hyperinflammatory to immunosuppressive immune responses, and often protracted failure to return to homeostasis (Cavaillon et al., 2020; van der Poll et al., 2017). Early organ dysfunction in sepsis is reversible and likely an adaptive strategy to withstand excessive inflammation and limit damage (Stanzani et al., 2019). A prolonged disruption of normal organ metabolism can however become maladaptive and eventually irreversible (Pool et al., 2018). Collectively, these factors and the systemic nature of sepsis introduce a highly complex pattern of overlapping molecular disease trajectories promoting organ damage through disparate processes such as pathological inflammation and immunosuppression, altered microcirculation and endothelial permeability, hypercoagulability, hypoxia, hypovolemia, metabolic disruption, and mitochondrial dysfunction (Lelubre and Vincent, 2018; Maslove et al., 2022).

Proteomics can provide novel insights into time-resolved organ- or cell-specific septic proteotypes, i.e. an assessment of the state of the proteome at any particular time or proteostates (Golden et al., 2021; Lapek et al., 2018; Toledo et al., 2019). Sepsis is reflected in organotypic proteostates during animal challenge models, where each organ displays unique signatures of highly enriched proteins generating organ-specific combinations of inflammatory, immunological and metabolic disease trajectories. Sepsis is also characterized by vascular leakage resulting in translocation of cells and blood plasma proteins into organs and a reciprocal translocation of tissue proteins into the blood plasma leading to a substantial increase in tissue-specific protein profiles in blood plasma (Malmstrom et al., 2016).

While criteria for diagnosing sepsis have been revised, standard protocols for therapy have remained largely unchanged for three decades due to the failure in developing new sepsis-specific therapies (Cavaillon et al., 2020; Marshall, 2014). Currently, prevention and management of septic organ dysfunction comprises four main strategies: pathogen elimination by antibiotics, stabilization of hemodynamics using fluids and vasopressors, organ system support, and judicious use of immunomodulatory therapies like glucocorticosteroids (Lelubre and Vincent, 2018). Over the last four decades, more than 120 clinical treatment trials have been conducted and all, expect one short lived example, have failed. An enormous degree of variability in sepsis presentation complicates delineation of patient subgroups/subphenotypes with similar clinical and molecular features (Cavaillon et al., 2020; Marshall, 2014; Scicluna et al., 2017; Seymour et al., 2019). Lacking adequate tools to objectively score the multi-faceted heterogeneity in septic organ failure, it remains highly challenging to evaluate new therapies and define optimal treatment windows. Furthermore, sepsis therapy has focused disproportionately on hyperinflammation due to lingering knowledge gaps in pathophysiology (Van Wyngene et al., 2018). Beyond activation of immune cells, coagulation, complement and cytokines by bacteria/bacterial products, recent reports suggest that metabolic deregulation due to aberrant mitochondrial pathways is another vital process driving septic organ dysfunction (Singer, 2014). Cessation of cellular functions without evidence of excessive cell death may be partly explained by loss of mitochondrial homeostasis and the ensuing bioenergetic disturbance (Takasu et al., 2013). The exact mechanisms of this dysfunction have not been elucidated in detail, hindering development of novel mitochondria resuscitating drugs as adjuvant therapies.

Failed intervention trials may also reflect our limited understanding of molecular treatment responses introduced by tested interventions in preclinical models. Commonly used readouts like survival and inflammatory markers offer little insight into the organ dysfunction status of experimental animals. More elaborate strategies to objectively quantify intervention efficacy at a molecular level, and to define on- and off-target intervention-specific effects, are needed Marked differences in organ- and cell-specific sepsis proteotypes suggest treatments exert a non-uniform effect across organ systems, but it remains unclear to what degree interventions can revert the thousands of dysregulated proteins and how such reversions would vary across organs. Furthermore, treatments may introduce unexpected changes unrelated to sepsis that may cloud the evaluation of treatment benefits for a given intervention.

Here we provide a strategy to define intervention-sensitive proteotypes objectively and quantitatively in animal infection models using a combination of conventional markers of sepsis, proteomics, and systems biology. Through a defined inoculum murine *Escherichia coli* intraperitoneal challenge model and time-resolved deep proteome maps of organ and blood compartments, we generate a previously unknown perspective of the *in vivo* molecular events preceding and associated with organ damage in sepsis. We then capitalize on this knowledge based to develop an unbiased and proteome-based strategy to score therapeutic effects of antibiotics and corticosteroids, revealing novel time-dependent, intervention-specific and synergistic effects in an organ- and blood compartment-specific manner during disease. We anticipate this approach will allow researchers to better characterize treatment effects in preclinical models of sepsis, define treatment-indicative biomarkers based on species-conserved regulatory networks that can be tracked in human studies, and improve the translation of novel sepsis therapies from preclinical models to humans.

## Results

### The multiorgan proteome landscape of murine sepsis and effects of molecular therapeutic interventions

To define the multiorgan landscape of sepsis and uncover molecular therapeutic intervention responses, we applied a combinatorial strategy based on pathophysiological read-outs such as cytokines, immune cells, organ damage, and bacterial burden in blood and organs, coupled to a deep sepsis proteome tissue atlas, thus describing the temporal proteome restructuring in plasma and key organs at high risk of developing damage (**Fig. 1A**) (Singer et al., 2016). C57BL/6J mice were inoculated intraperitoneally (IP) with 10^4^ colony forming units (CFU) of *E. coli* strain O18:K1 and the time-resolved systemic molecular sepsis response 0, 6, 12 and, 18 hours post-infection (h.p.i.; n=23) determined to identify possible therapeutic treatment windows (**Fig 1A-C)**.

**Figure-1.**
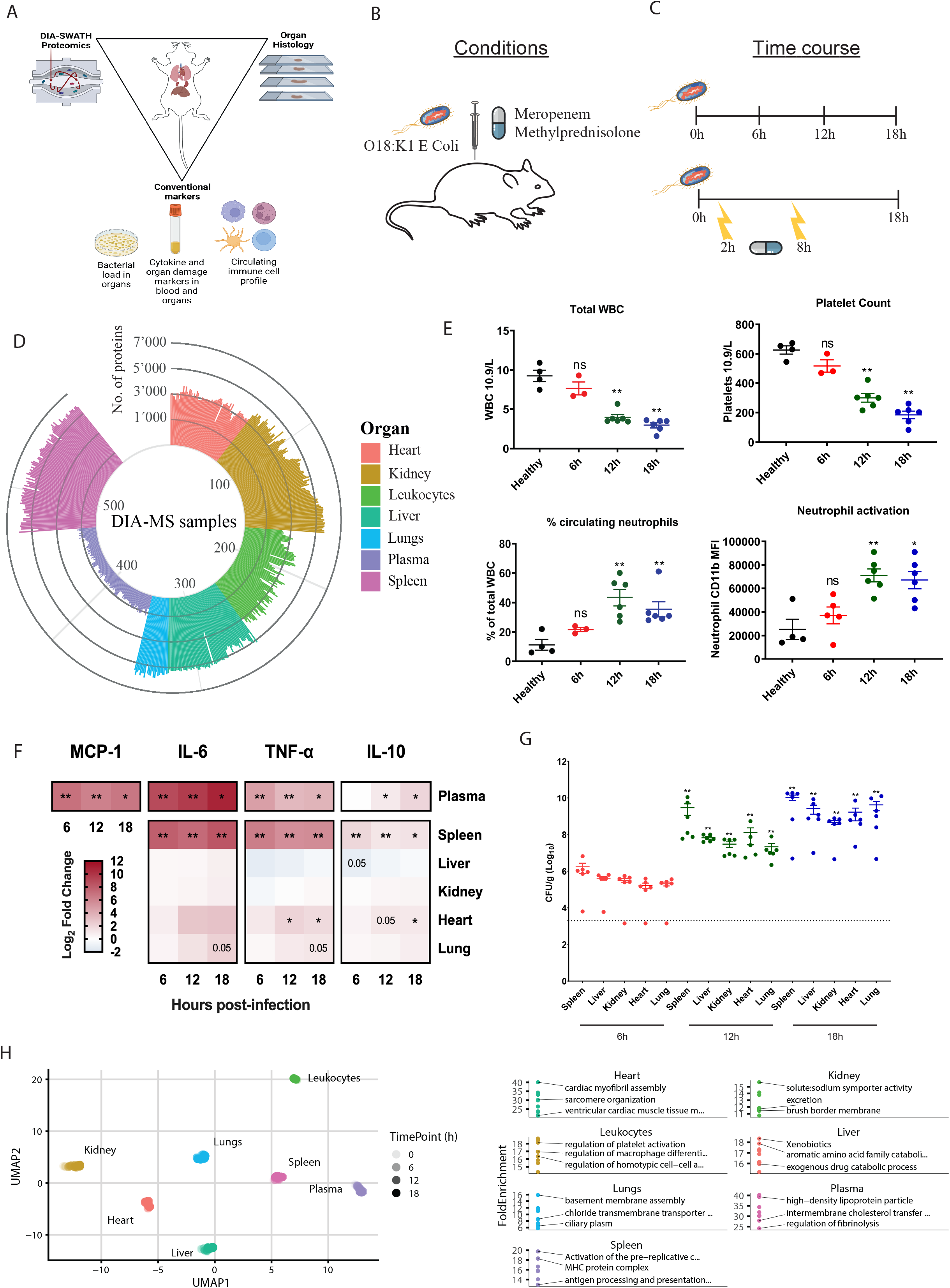
Proteomic landscape of murine sepsis. A) Model of sepsis and interventions. C57BL/6J were infected with *Escherichia coli* O18:K1 strain. B) Time course of infection and intervention. C57BL/6J were infected with 10^4^ *E. coli* and C) organs were harvested at 0, 6, 12 and 18 h.p.i. (n=6/timepoint) and analyzed with DIA-SWATH maps, organ histology and conventional markers like bacterial load in organs, cytokine profile, organ damage markers and circulating immune cell profile. For the intervention study, infected animals were treated with 30 mg/kg methylprednisolone and 10 mg/kg meropenem (n=4/treatment). D) Total split of the samples and detected proteins using DIA-SWATH MS. E) Leukocyte pool during sepsis time-course. Total white blood cell count (10^9^/L), blood platelet count (10^9^/L), percentage of circulating neutrophils (% of total WBC) and median fluorescence intensity (MFI) of activation marker CD11b on neutrophils as determined by flow cytometry. F) Cytokine levels in plasma and organ homogenates depicted as log_2_ fold-change. G) Viable bacterial load in organs. Viability was assessed in spleen, kidney, lung, liver, heart and spleen. Colony forming units (CFU) expressed as CFU/g of tissue (See Figure S2 for all combined physiological data). H) UMAP projection of organs over time. UMAP projection depicting organs segregation over disease progression. The table indicates the functional hallmark groups associated with each organ.

We then repeated the IP *E. coli* inoculation followed by administration of 10 mg/kg meropenem or 30 mg/kg methylprednisolone at 2 or 8 h.p.i. Meropenem is a broad-spectrum bactericidal β-lactam antibiotic used in empiric sepsis therapy but with limited knowledge of its specific effects on the host proteome response. Methylprednisolone is a synthetic glucocorticoid prescribed for its anti-inflammatory and/or immunosuppressive effects, and frequently studied as a sepsis intervention without consistent benefits (Annane et al., 2009). We further studied combinations of the two treatments at 2 and 8 h.p.i. to examine potential synergistic, antagonistic, or orthogonal molecular intervention effects.

In each experiment, we harvested heart, kidney, liver, lung, and spleen along with the blood compartment comprised of the leukocyte pool and blood plasma to construct a murine sepsis spectral library from fractionated organs and cell lysates. This library contained 4,614,190 peptide spectrum matches covering 102,829 peptides matching to 11,510 proteins (**Figure S1**). We then constructed 557 DIA-MS proteome maps from the harvested organs, cells and plasma from each experiment, and used the corresponding spectral library to extract peptide and protein quantities, identifying 9620 unique protein groups; all data, including spectral library, 1500 proteome maps and result files are available from Proteome exchange (**Fig. 1D**) (under accession code XXXXX). To the best of our knowledge, the resource presented here provides the largest compendium of label-free DIA-SWATH maps for an animal model of infection.

### Multi-organ time-dependent monitoring of murine E. coli sepsis

Over the tested time course, animals progressively developed severe sepsis characterized by a time-dependent reduction in weight (**Figure S2**), total white blood cell counts, and platelets, coupled to a reciprocal increase in activated CD11b^+^ neutrophils (**Fig 1E**). A significant increase in inflammatory cytokines was identified in spleen and plasma, and in the heart at later time points (**Fig 1F)**. The spleen had the highest bacterial colony forming unit (CFU) counts during early stages of the infection (t = 6 h.p.i) (**Fig 1G**). Although the proteome data segregated the individual organs by response to sepsis, their intrinsic tissue proteomes remained strongly distinct and enriched for typical organotypic protein machineries, such as cardiac myofibrils in the heart, high-density lipoprotein particles in plasma, and antigen processing in the spleen (**Fig, 1H**).

### Profiling septic organ damage and tissue protein leakage in plasma

To verify that infected animals developed multiorgan failure, we used a combinatorial strategy to assess organ damage markers, histology and proteomics, plasma concentration of certain proteins such as acute phase proteins, immune cell derived proteins, and cytokines. The observed changes were organotypic and characterized by an increase in splenic white pulp in after 6 h, reduction of hepatocyte nuclei, time-dependent increase of mesangial proliferation, and fibrin clot formation in all organs (**Fig 2A-B**). At later time points, established organ damage markers such as lactate dehydrogenase (LDH), blood urea nitrogen (BUN), alanine aminotransferase (ALT), and cardiac troponin T were significantly elevated in plasma (**Figure S2)**. The manifestations of organ damage and inflammation were supported by proteome data, which revealed a time-dependent increase of protein groups related to fibrin clot formation, neutrophil degranulation, interferon-beta (IFN-β) response, and platelet degranulation in plasma and organs, all of which in excess are linked to organ dysfunction (Hotchkiss et al., 2016) (**Fig 2C**). These increases in established organ damage markers and inflammatory proteins were accompanied by a marked change in blood plasma proteome homeostasis, with a marked rise of organ/cell resident proteins entering the circulation (**Fig. 2D**). In total, 275 tissue-abundant and tissue-specific proteins increased in plasma in a time-dependent fashion (**Fig. 2E**), including those normally involved in specialized tissue functions such as cell adhesion, glycolysis, gluconeogenesis, glutathione metabolism, and aldehyde metabolic process (**Fig. 2F**). Several functional groups of leaked proteins were overrepresented in specific organs, such as α-amino acid metabolic process in the liver, aldehyde metabolic process in the lungs, and glutathione metabolism in the heart and the liver (**Fig. 2G**). In contrast to conventional organ damage markers, over 37% (n=103) of the leaked proteins increased organ abundance during sepsis, paralleling their release into the plasma during organ damage **(Fig. 2F)**. Organ damage-associated tissue leakage occurs later at 12h and 18h and is rather stochastic, likely due to protein localization and abundance, ease of release into plasma, and the degree of organ damage. Liver is the major contributor to this signal likely due to its size, vascularization, and significant baseline contribution to the plasma proteome. Our results perhaps highlight an underappreciated magnitude of leakage of functionally connected tissue proteins into blood plasma where they are normally absent and can serve as a future resource to define novel plasma-based organ damage markers and delineate distinct damage responses.

**Figure-2.**
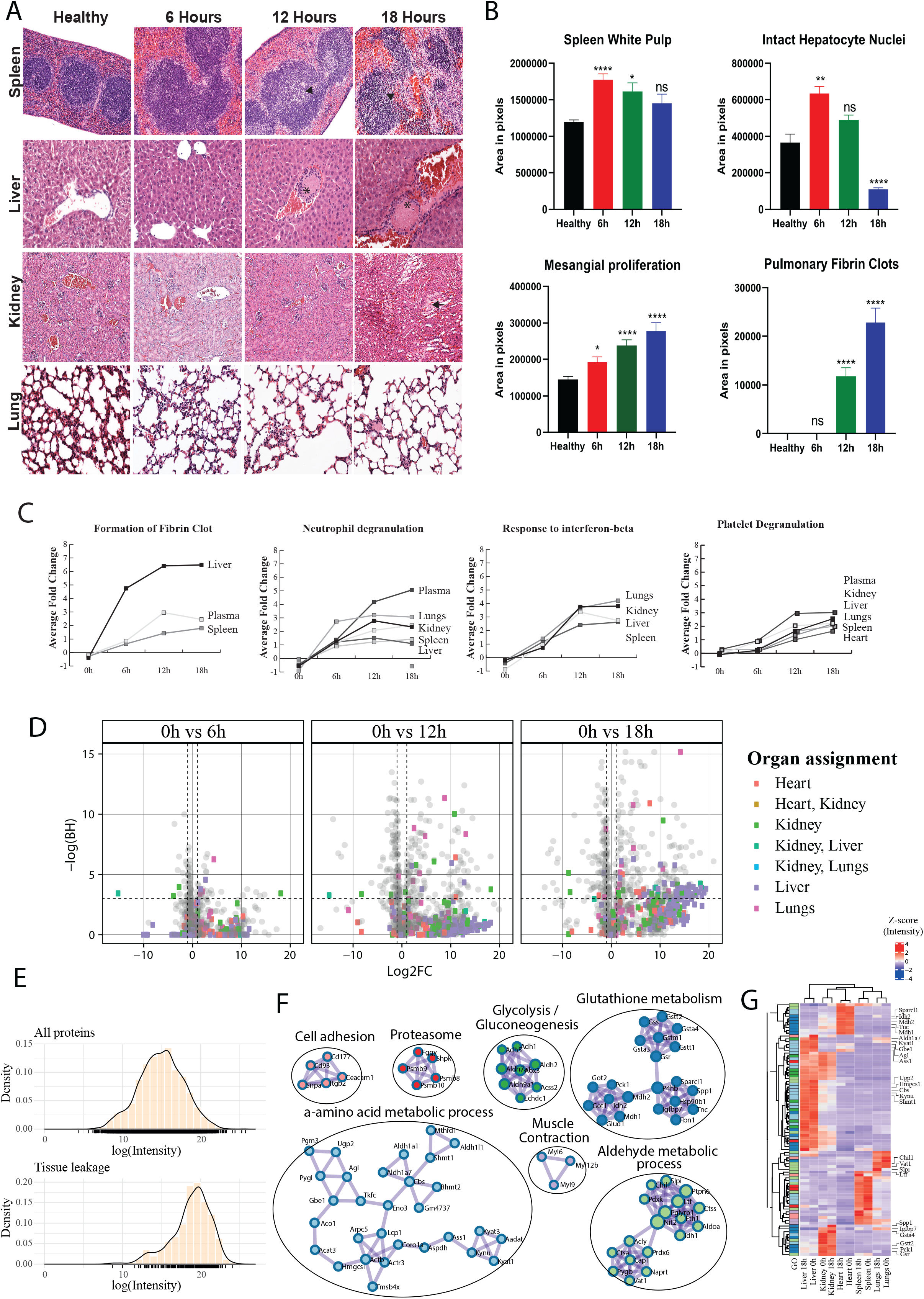
Organ damage during sepsis. A) Histology of organ damage. Representative hematoxylin and eosin (H&E) staining of spleen, liver, kidney and lung. The images depict focal necrosis in the white pulp of spleen (black arrowhead), appearance of clots in the liver (asterisk), tubular necrosis in the proximal tubules in kidney (black arrow) and appearance of clots in the alveoli in lungs (white arrow). B) Semi-automatic quantification of H&E staining. Images were quantified with FIJI. All conditions were compared with healthy (0h) using Kruskal-Wallis with *** p<0.0005, ** p<0.005, * p<0.05 and ns = non-significant (See Figure S2 for a combined panel of all physiological data). C) Average increase of inflammatory components like formation of fibrin clot, neutrophil degranulation, response to interferon-beta and platelet degranulation across organs and plasma. D) Appearance of tissue leakage proteins in blood plasma. Volcano plots showing the contributions of various organs to the tissue leakage protein pool. Cut-off for Log2FC = +/1.5, -logp = +/-4. Colors indicate predicted organ assignments. The proteins were assigned to an organ if they were at least >20 fold more abundant in one organ with a corrected p-value > 0.05. E) Density plot of log organ protein intensities of all proteins and for the tissue-specific proteins identified in plasma. F) Functional enrichment of leaked proteins. G) Heatmap showing normalized intensity of the tissue leakage proteins in the organs under uninfected state and after 18 h.p.i. The colors in the first column indicates the functional groups shown in F.

### Temporal profile of organotypic proteome responses in E. coli sepsis

The observed dysfunction across individual organs is likely to disrupt intricate crosstalk essential for maintaining homeostasis (Bauer et al., 2018; Caraballo and Jaimes, 2019). Although individual organ proteomes segregate over time in an organ-dependent manner during sepsis (**Fig. 3A**), we lack a basic comparative understanding of organ dysfunction between these organs from infection to severe disease (Caraballo and Jaimes, 2019), including differences in the onset and rate of decline. We therefore performed pairwise differential protein abundance testing to determine when differentially abundant proteins (DAPs) arose and if they persisted with increasing disease severity. Over the sepsis course, a time-dependent increase in the number of DAPs was observed (**Fig. 3B**), starting to plateau around 12 h (**Figure S3a**), indicating a breakpoint between 6-12 h where the increase in response is no longer linear. The pattern and magnitude of DAPs subdivided organs into two groups that we refer to as responder organs (liver, leukocytes, and spleen) or bystander organs (lung, heart, kidney, and plasma) (**Fig 3B** and **Figure S3a**). This subdivision may be connected to the known activation of the leukocyte population (Fig 1E, CD11b^+^) and changes in metabolisms during sepsis.

**Figure-3.**
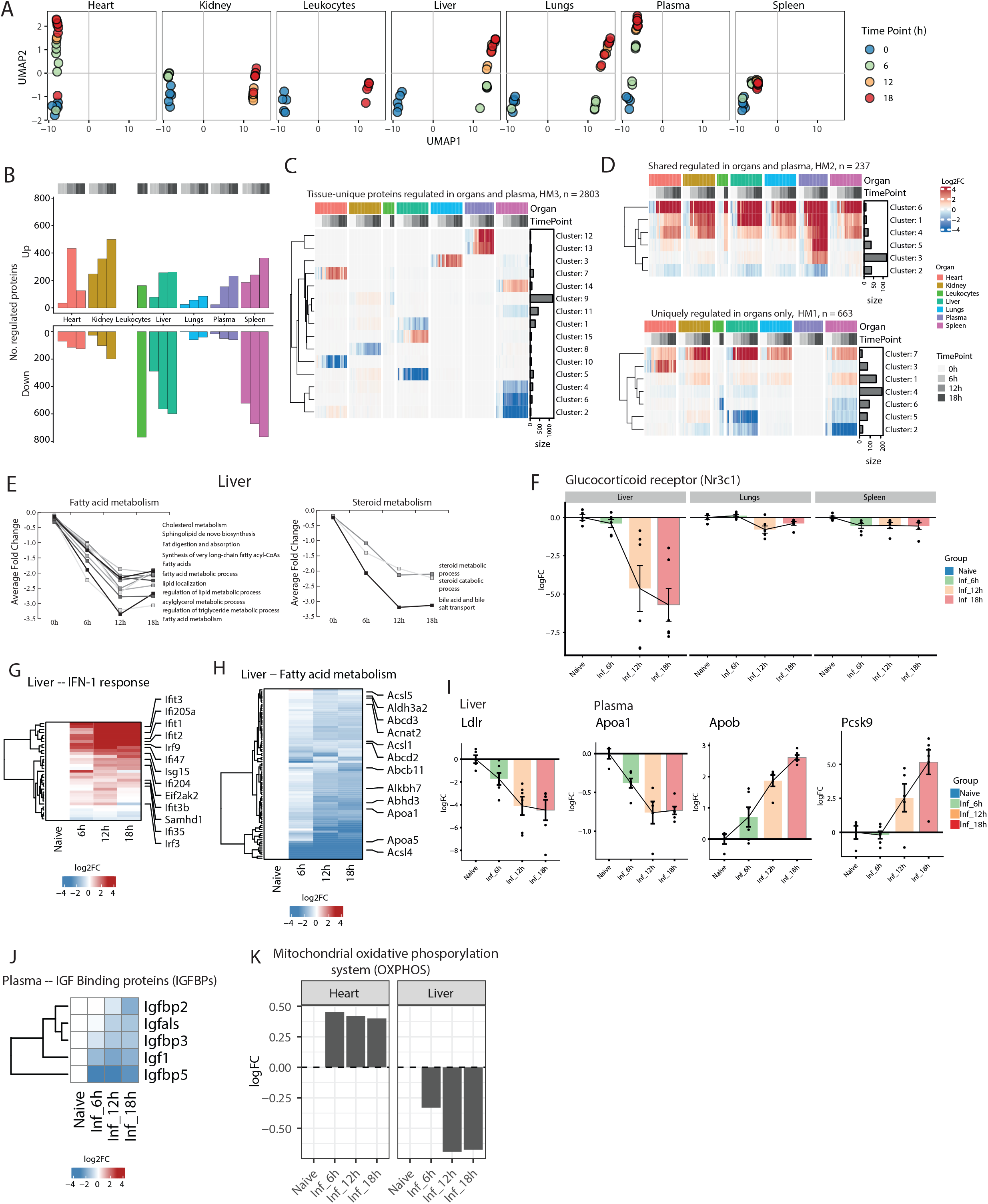
Temporal profile of organotypic proteome responses in E. coli sepsis. A) UMAP projections of segregated organs over time. B) The number of differentially abundant proteins (DAP) across organs and the blood compartment over time. C) Heatmap showing average normalized intensity of protein clusters that were significantly more abundant on one organ. D) Upper heatmap in the panel shows protein cluster with significant increased protein abundances in plasma in at least one organ. Lower heatmap in the panel shows protein clusters with significant increased protein abundances in at least one organ but not in plasma. E) Organotypic responses in liver depicting significantly lower levels of proteins associated with functional groups involved in fatty acid and steroid metabolism. F) Time-dependent changes in abundance level of the glucocorticoid receptor (Nr3c1) in liver, lungs and spleen. G) Heatmap showing the increased levels of type-1 interferon (IFN-1) response proteins over time in liver. H) Heatmap showing reduced levels of fatty acid metabolism proteins over time in liver. I) Time-dependent changes in abundance level regulating apolipoproteins in plasma and liver. J) insulin-like growth factor (IGF-1) signaling in plasma over time and the. K) mitochondrial oxidative phosphorylation (OXPHOS) components in heart and liver over time.

The proteome response can be broadly divided into three categories: DAPs that are uniquely regulated in one organ or plasma, regulated DAPs shared across organs, and uniquely plasma and tissue regulated DAPs. Clustering the DAPs revealed that all organs including plasma were exposed to a synchronized time-dependent increase in fibrin clot formation and neutrophil degranulation (**Fig. 2C, 3C-D and Figure S3b**). Response to IFN-β was observed only in organ tissues but not in blood plasma. Other systemic proteome changes confined to organs and tissues but not plasma included functions like altered Toll-like receptor-4 pattern recognition receptor signalling pathway, regulation of the insulin pathway, and apoptosis (**Figure S3b**). Apart from this, bystander organs were relatively refractory to intrinsic organotypic alterations, as the above-mentioned changes represented a notable proportion of the observed response (**Figure S3c**). In contrast, responder organs were characterized by a substantially higher number of sepsis-reduced DAPs. Distinct changes in organotypic biological processes include steroid and fatty acid metabolism in liver, altered glucose metabolism and changes to the lipoprotein particle composition in plasma, and reduction of ribosome levels in heart (**Fig. 3E, Figure S3b** and **Figure S3d**). In addition, we observed time-dependent changes in several hallmark proteins studied in human sepsis such as a reduction in liver glucocorticoid receptor (GR)-associated pathways in the liver such as IFN-1 signalling (Flammer et al., 2010), fatty acid metabolism and gluconeogenesis (Van Wyngene et al., 2018) (**Fig. 3G-H**). The low-density lipoprotein receptor (LDLR) was also reduced in liver accompanied by increased levels of proprotein convertase subtilisin/kexin type 9 (PCSK9) in plasma, culminating in altered balances between the high-density lipoprotein-associated apolipoprotein a1 (APOA1) and low-density lipoprotein-associated apolipoprotein b (APOB) (**Fig. 3I**). Concurrently, over the time course, a drop in IGF-1 and insulin-like growth factor binding proteins (IGFBPs) was noted, increasing bioavailability of IGF-1 in plasma (**Fig. 3J**), with only minor changes in the mitochondrial oxidative phosphorylation system (OXPHOS) (**Fig. 3K**).

In conclusion, sepsis instigates multiple organ failure and a massive and organotypic proteome reorganization of all analysed organs in this model. The temporal profiles demonstrate that many responses increase linearly during the early phase of the disease to plateau during the later stages, raising the possibility that early stages represent treatment sensitive proteostates.

### Definition of proteome intervention responses during sepsis

To test treatment responses, we administered early or late glucocorticoid (Gcc) or meropenem (Mem), or combinations of the two, at 2 or 8 h.p.i. (**Fig**. **1A** and **4A**). At 2 h.p.i., the infection was systemic but with low organ CFUs (10^4^-10^5^ CFU/g of tissue) and a decrease in weight (**Figure S4a**). By 8 h.p.i., the animals were progressively sicker with a 10-to 100-fold increase in organ CFU and an additional 8% decrease in weight (**Figure S4a**). The 8 h.p.i timepoint falls within the time-range before the proteome response starts to plateau (**Fig. 3**). Mem intervention eliminated *E. coli* CFU in the heart, lung, and kidney, with low residual CFU in the spleen and liver independent of intervention time (**Fig. 4B, Figure S4b**). Conversely, early administration of Gcc increased *E. coli* CFU in all organs compared to untreated animals. Interestingly, early administration of Gcc also blunted bacterial killing by Mem, possibly by decreasing the levels of NADPH oxidase component cytochrome B-245 Beta Chain (CYBB) in the leukocyte population required for effective oxidative burst (Rae et al., 1998) (**Figure S4a)**. In contrast, late combination intervention with Gcc and Mem resulted in a markedly reduced bacterial load in the organs, altered levels of proinflammatory cytokines in the plasma and spleen, sepsis-induced weight loss, and increased WBC levels, together indicating that Gcc therapy accentuates some of the positive effects exerted by Mem (**Fig 4B, Figure S4b**).

**Figure-4.**
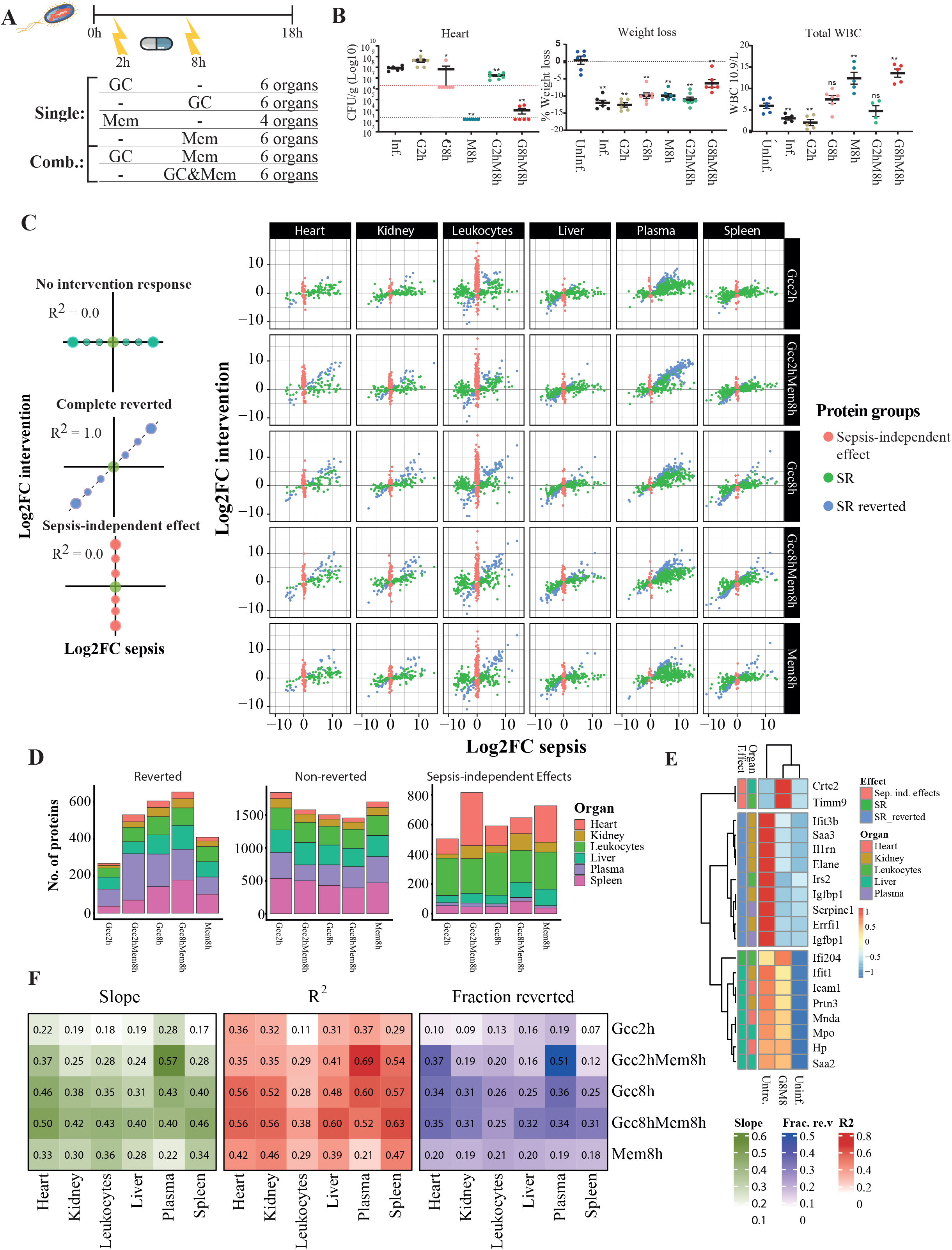
Scoring intervention effects in organs and the blood compartment. A) Scheme of infection of C57BL/6J and treatment intervention administration. B) Physiological markers. Bacterial load in heart (CFU/g of tissue), weight loss (%weight loss prior to infection) and total white blood cell count expressed as 10^9^/L. All animals were harvested at 18 h (n=6 per group) C) Scheme and scatter plots depicting the three categories of proteome regulation by treatments: reverted, non-reverted, and side effects across organs and the blood compartment. The colors indicate three protein categories referred to as the reverted (blue), non-reverted (green) and sepsis-independent effects (SIE)(red) D) Stacked bar plots showing the number of regulated proteins for interventions in different organs in the reverted, non-reverted and SIE categories. E) Heatmap of hallmark proteins showcasing the effect of combined glucocorticoids and meropenem administered at 8 h (GccMem8h) comparing infected and healthy animals. The first column in the heatmap shows the scoring categories by color. F) Heatmaps showing the slope, R^2^ and fraction of reverted proteins (of total) for the different interventions across different organs.

The proteome response profiles supported the above findings. Extensive molecular disturbances instigated by sepsis in our model were complex and acute, and despite systemic effects of the interventions, both glucocorticoids and antibiotics could only partially revert the proteome to resemble the uninfected state. Plotting the proteome intervention response both as fold-change in sepsis vs. untreated (“Sepsis Response”) and fold-change intervention vs. sepsis (“Sepsis Response Reverted”) segregated the response into three categories (**Fig 4C)**. One category represents proteins we designate sepsis response proteins, changed by sepsis but unaffected by the interventions (green; no intervention response). The second category is comprised of proteins that were changed by sepsis but significantly reverted by the interventions towards the uninfected baseline (blue; complete reverted), designated sepsis reverted. A third category included proteins that did not change due to sepsis but only by the treatments themselves, designated sepsis-independent treatment effects (red; sepsis independent) (**Fig 4C and D**). On this aggregated organ-proteome level, the non-reverted sepsis signal was surprisingly strong indicating that the tested interventions can only partly counteract the sepsis-induced pathways (**Figure S4c**).

Of the tested interventions, Gcc8hMem8h (glucocorticoid plus meropenem each at 8 h) significantly reverted the highest number of proteins, and to a large degree dampened the sepsis proteome response of the non-significantly reverted proteins (**Fig 4D** and **Fig. 4E**). To quantify the intervention impact, we calculated the slope and R^2^ value (**Figure S4d**) and the percentage of the reverted sepsis response for the proteins in the two protein categories for all organ-intervention combinations (**Fig 4F**). In this calculation, a slope and R^2^ of 1.0 equals a complete reversion of the sepsis response by the interventions. These calculations show that the late combination of Gcc8hMem8h resulted in the highest average R^2^ (0.54), the highest slope (0.44) and the largest fraction of reverted proteins from the sepsis response (31%), with the strongest beneficial treatment effect on the heart and the leukocyte pool. Gcc2h alone represented the opposite extreme, with only minimal reversion of the sepsis response (12%, R^2^=0.29, slope=0.17) possibly due high bacterial CFU. This effect was most noticeable in the kidney and spleen, where early Gcc2h resulted reverted less than 10% of the sepsis-induced proteins. One strong reversion effect of Gcc2h however was a significant reduction in the response to IFN-β (**Figure S4c**). To benchmark the magnitude of the quantifiable intervention response, we repeated the experiment including Mem2h as a positive control. Mem2h had a substantially higher average R^2^ value (0.86), and slope (0.77) compared to the other treatments and reverted on average close to 80% of the sepsis significant DAPs, highlighting the importance of early treatment (**Figure S4d** and **Figure S4e**). Also in this experiment, Gcc8hMem8h resulted in a higher R^2^ (0.65) and percentage of reverted proteins (31%) on average compared to the other treatments (**Figure S4e**).

These results show that the changes in the proteome landscape can be used to quantify the intervention-impact at an organ level. Intervention effects ranged from 7% reverted proteins with an R^2^ value of 0.11 to a maximum effect of 85% reverted protein and a R^2^ value of 0.92. In addition, our analysis revealed a third and more unexpected category of proteins that were unaltered during sepsis but significantly changed because of the interventions during septic conditions: sepsis-independent effects (**Fig. 4C**-**D**). Proteins in this category are easily overlooked yet likely represent an important parameter in assessing interventions effects, especially as the number of proteins in this category was surprisingly high (>600 proteins), identified most frequent in the heart and the leukocyte pool **Fig. 4D**).

### Scoring of intervention altered pathways

To further explore the functional effects among the reverted proteins, we applied the same proteome scoring shown in Figure 4 to enriched functional protein categories. This analysis revealed that the late interventions had strong organotypic effects on the altered inflammatory (**Fig. 5A**) and metabolic responses (**Fig. 5B**). Reversion of inflammatory responses was most pronounced in the kidneys, with responses such as regulation of cytokine production, cell killing, regulation of immune effector processes, and carbohydrate metabolic processes (Toledo et al., 2019) were strongly reverted back towards the uninfected state by the late treatments (**Fig. 5A and Figure S5a**). The combinatorial effect of Gcc8h and Mem8h potentiated the effects exerted by the mono-interventions in the kidney (**Fig. 5C**), with Enrichment analysis against the TRRUST database in the kidneys showing that Gcc8hMem8h specifically impacted the pro-transcriptional regulators of inflammation like Nfkb1 (p = 10^−3^), Jun, (p = 10^−2.4^), and Sp1 (p = 10^−2.2^) (Kusnadi et al., 2019) (**Figure S4b**). These inflammation-related functional categories were reverted in the heart to a lesser extent, with no noticeable intervention effect in the spleen or liver.

**Figure-5.**
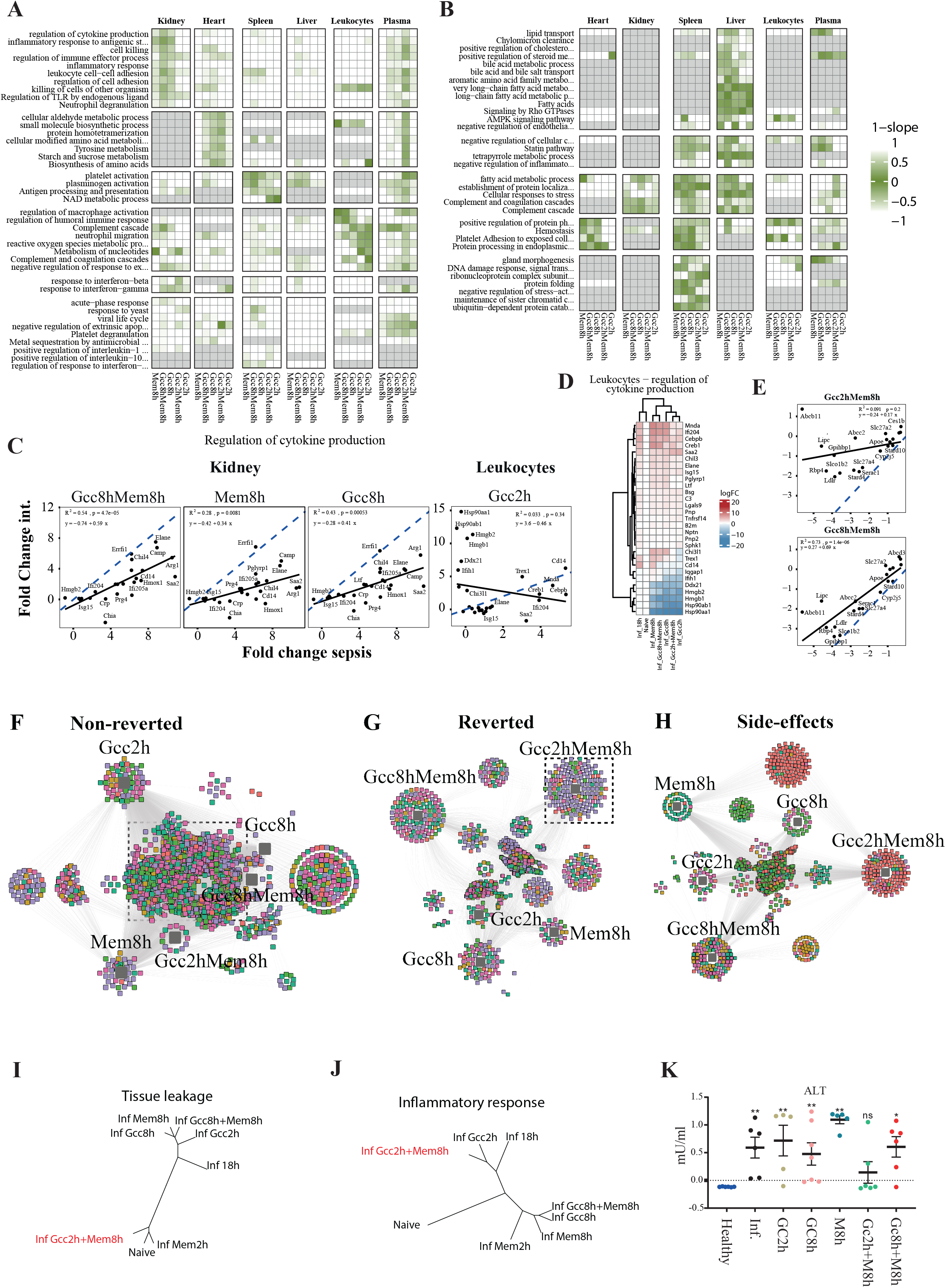
Organotypic intervention response networks in sepsis. The members of the three protein categories shown in **Figure 4** were subjected to functional enrichment using Metascape (Zhou et al., 2019). A 1-slope value was calculated for each protein category, where protein categories close to 0 are colored indicating the intervention effect per treatment group shown as a heatmap for functional groups associated with increased protein abundance (A) and functional groups with decreased protein abundance (B). C) Example scatter plots of fold-change sepsis vs fold-change intervention used to calculate the values in (A-B) for protein members in the functional group ‘regulation of cytokine production’. Solid line shows the linear regression line, and the dotted line shows a slope of 1, which indicates the theoretical complete reversion all the protein members back to uninfected state. D) Heatmap showing the normalized protein intensities after intervention for the proteins associated with ‘regulation of cytokine production in leukocytes. E) Scatter plot of fold-change sepsis vs fold-change intervention for the protein members in the liver for the functional category ‘lipid transport’ in animals treated with Gcc2hMem8h or Gcc8hMem8h. The solid line indicates the linear regression line, and the dotted line shows a slope of 1 for comparative reasons. F-H) The member of the three protein categories related to their respective intervention using the p-value as edge (cut-off was corrected p-value of >0.05) and visualized using cytoscape for non-reverted (F), and box with dashed lines represents shared proteins for all interventions, G) reverted (dashed box contains the reverted proteins with Gcc2hMem8h) and H) sepsis-independent effect (SIE) by interventions across different tissues. I) Total tissue leakage proteins into plasma by different interventions. Colors indicate organ associations. J) Treatment trees for known inflammatory proteins and the 275 tissue-leakage proteins in plasma for different interventions. K) Plasma levels of alanine aminotransferase for different interventions expressed as mU/ml.

We measured cytokines such as IL-6, TNF-α and MCP-1 in plasma using immunoassays and found their levels reduced by all interventions except early Gcc (**Figure S4b**). A similar trend was seen for IL-6 reduction in the spleen, which produces the cytokine during endotoxemia (Park et al., 2014), reduced except for early Gcc. Organ proteomes also revealed early administration of Gcc did not alter the systemic inflammatory response in any organ including the kidney. In the blood circulation, GccMem8h elevated leukocytes and platelets, while reducing the number of activated CD11b^+^ neutrophils, indicating a reversal of disease (Zhou et al., 2005); conversely, early Gcc treatments were associated with leukopenia and thrombocytopenia. The early Gcc intervention significantly increased concentrations of innate immune response proteins induced in sepsis, such as high mobility group proteins (HMGB-1, −2) (Yanai et al., 2011) and the interferon-type 1 pathway (Ifih1, ifi204) (Lei et al., 2021). These pathways regulate IFN-β responses (Lei et al., 2021), participate in immune recognition of damage-associated nucleic acids as well as heat shock proteins (HSP90AA1, HSP90AB1) (Wan et al., 2020), and are involved in activation of interferon regulatory factors −1 and −3 (IRF1, 3) and STAT1(Lei et al., 2021; Puthia et al., 2016) (**Fig. 5D**). In addition, early Gcc administration also reverted proteins like CEBPB, a transcription factor regulating the expression of genes involved in immune and inflammatory responses, and CREB1, which is a phosphorylation-dependent transcription factor that stimulates transcription upon binding to the DNA cAMP response element (CRE) (Ruffell et al., 2009).

The host-response to bacteria in our model showed a marked elevated inflammatory response in all organs, an organotypic reduction of DNA repair and transcription in the spleen, and fatty acid, steroid, and retinol metabolism in the liver (**Fig. 3**). Several of these processes were distinctly reverted by the late interventions, underscoring the importance of timing when administering interventions (**Fig. 5B**). Most apparent was the partial reversion of the metabolic module in the liver including lipid transport, regulation of cholesterol and steroid metabolism, long-chain fatty acid metabolism, and bile acid metabolic processes. The transcriptional regulatory network connected to this effect was HNF1 homeobox A (p = 10^−2.6^), which is expressed highly in the liver and regulates expression of several liver-specific genes (Courtois et al., 1987). Repression of lipid metabolic pathways in the liver and elevated lipid metabolites are associated with Gram-negative sepsis (Van Wyngene et al., 2020). Apart from its primary metabolic function, lipid transport also facilitates clearance of toxic lipopolysaccharide (LPS) through high-density lipoprotein (HDL) particles from circulation in the liver (Walley et al., 2015). One of the strongest intervention-reverted proteins in this group was LDLR. Reduced PCSK9 function is associated with increased pathogen lipid (LPS) clearance via the LDLR, a decreased inflammatory response, and improved septic shock outcome (Walley et al., 2014). In support of these earlier observations, our data indicate repression of lipid transport proteins in the liver of septic animals **(Fig. 5E**).

### Organotypic intervention response networks in sepsis

To further investigate the proteins that remained unaltered by the treatments and to determine if interventions specifically alter the host response, we plotted drug response networks. In these networks, significant intervention-protein associations within the three protein categories were connected with edges and organized into a network (**Fig. 5F-H**). The graphs show that over 1092 associated protein remained statistically non-reverted after all interventions ranging from 39 proteins in heart to 331 proteins in spleen (**box in Figure 5F**). Functional enrichment shows that there is a substantial degree of inflammation, such as neutrophil degranulation (p = 10^−37^) and complement and coagulation cascades (p = 10^−29^), that could not be reverted by any of the tested interventions. Interestingly, many of these functional categories were unaltered even after intervention with Mem2h, indicating an intervention refractory response that is triggered early in the sepsis response and that is maintained throughout the course of the disease (**Figure S5a**).

In general, Gcc2hMem8h reverted sepsis-induced pathways only to a minor extent (**Fig. 5A-B**). A noticeable exception to this was the specific and significant decrease of 92 sepsis-induced tissue leakage proteins in the plasma (**Fig. 5A and 5G**). These tissue leakage proteins are intracellular proteins statistically enriched from the liver (p = 10^−19^), that leak out into plasma during sepsis and are involved in biological functions such metabolism of amino acids and derivatives (p = 10^−51^), amino acid metabolism (p = 10^−35^), and carbon metabolism (p = 10^−34^) (**Figure S5b)**. To investigate if Gcc2hMem8h introduced a general reduction in the leakage of tissue proteins into plasma, we plotted intervention trees of the 275 tissue-abundant and tissue-specific that were elevated significantly during sepsis (identified in **Figure 2**). These intervention trees show that Gcc2hMem8h cluster with the uninfected animals and the gold standard treatment Mem2h **(Fig. 5I)**. In contrast, the increase in plasma inflammatory proteins were unaffected by Gcc2hMem8h where this intervention clusters with the untreated animals (**Fig. 5J**). As this intervention by itself does not completely clear bacterial CFU and does not decrease the inflammatory profiles or reduced organ damage, our results imply that Gcc decreases vascular permeability as indicated by the reduced levels of the organ damage marker ALT (**Fig. 5K**). These findings are relevant because they show that interventions can alter the levels of plasma biomarkers and organ damage markers without affecting inflammation and organ damage.

### Characterization of intervention-induced sepsis independent effects and mitochondrial dysfunction

In addition to the reverted/non-reverted proteins groups, we identified a group of proteins that did not change due to sepsis but only by the treatments themselves, designated sepsis-independent treatment effects. Most sepsis-independent response DAPs were downregulated at the organismal level by all interventions. These DAPs were strongly linked to mitochondria and were enriched for electron transport chain, mitochondrial transport, membrane organization, and translation (**Figure S4c**). Glucocorticoids and beta-lactam antibiotcs can influence mitochondrial metabolism (Scheller and Sekeris, 2003) and induce stress responses (Kalghatgi et al., 2013). But their individual/combined effects on mitochondria have not been described in sepsis. A pronounced presence of sepsis-independent effects DAPs was seen in the heart, especially DAPs that were downregulated (>300 proteins) (**Fig. 6A**). Many functional protein categories in the downregulated sepsis-independent effects response were mitochondria-related (**Figure S6a & S6b**). The two Gcc treatments lowered sepsis-independent effects DAPs (Gcc2h, 79 proteins and Gcc8h,108 proteins). In contrast, conditions where Mem was added late (Gcc2hMem8h, 315 proteins and Mem8h, 215 proteins) had higher sepsis-independent effects DAPs. Importantly, late combination treatment with Gcc8hMem8h (70 proteins) counteracted sepsis-independent effects DAPs introduced by Mem in the heart such as those involved in β-oxidation (**Fig 6B**). Their reduction in the heart during septic conditions by antibiotics has not been described to the best of our knowledge, whereas the reduction of cell redox homeostasis is known (Kalghatgi et al., 2013). In line with previous reports demonstrating cardioprotective effects of Gcc (Varvarousi et al., 2014), our data indicate that Mem in the setting of early Gcc or absent Gcc is associated with cardiac mitochondrial dysfunction.

**Figure-6.**
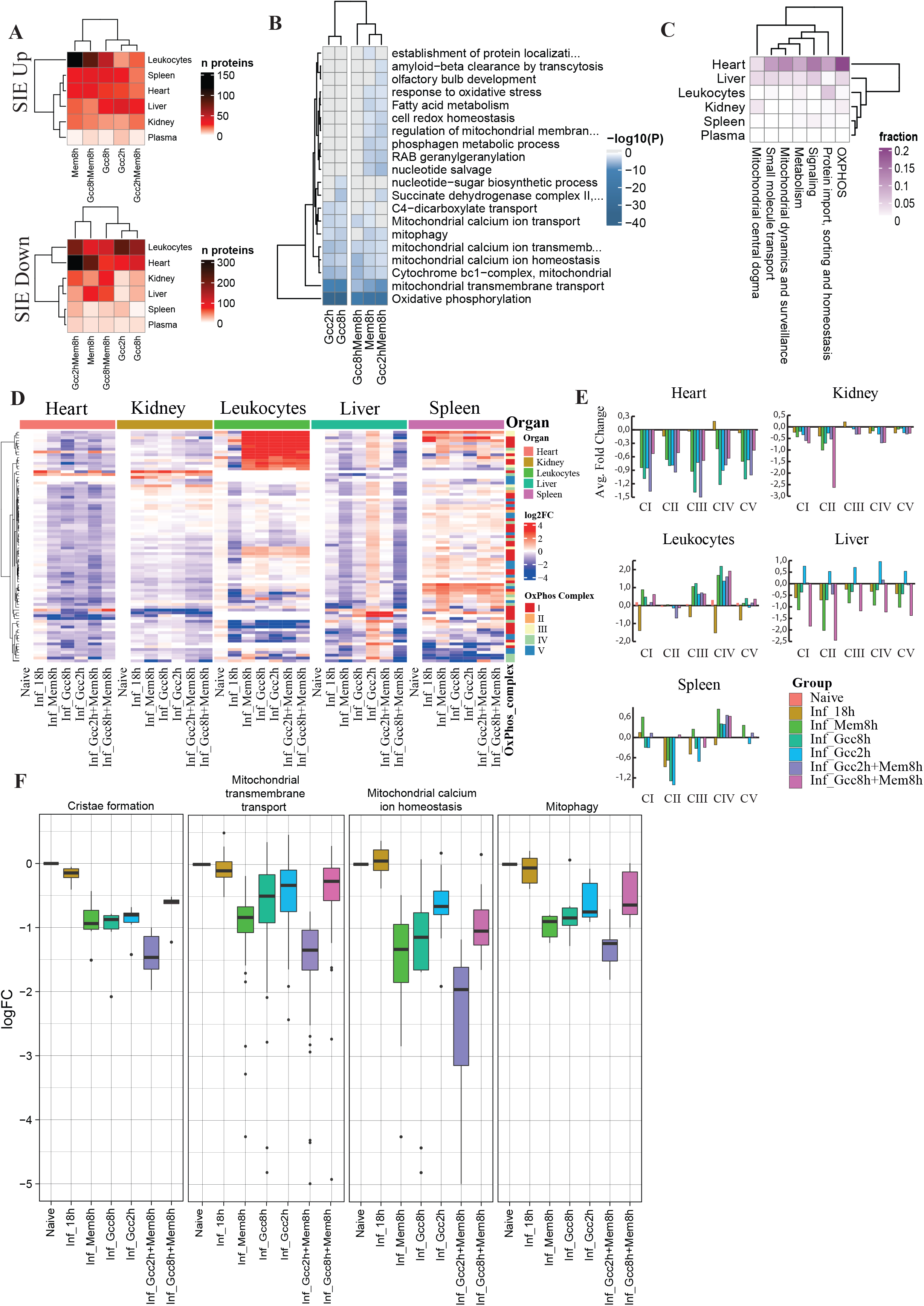
Sepsis-independent effect proteins, enrichment and dysfunction of cardiac mitochondria. A) The number of sepsis-independent effect (SIE) proteins from **Figure** 4 per organ and blood compartment separated into reduced or increased abundance levels. B) Functionally enriched protein categories of the SIE proteins in the heart. Color gradient indicates -log10(P). C) Mitocarta enriched terms of SIE proteins in organs and the blood compartment. D) Heatmap showing normalized proteins intensities for the proteins part of oxidative phosphorylation (OXPHOS) proteins in different organs. The color column indicates OxPhos complex I-V E) Fold-change of the average MS intensities for the OXPHOS complexes in organs. F) Box plots of the average MS intensities for the protein members in the mitochondrial terms associated with ‘cristae formation’, ‘membrane transport’, ‘calcium ion homeostasis’ and ‘mitophagy’. Colors indicate type of intervention.

Because of the pronounced enrichment of mitochondrial proteins in the sepsis-independent effects group, we selected all proteins annotated as mitochondrial proteins from MitoCarta 3.0 (Rath et al., 2021). This yielded 945 quantified mitochondrial proteins across the organs out of the 1140 catalogued proteins in MitoCarta, which we plotted based on the three protein categories shown in **Figure 6C** (**Figure S6c**). Functional enrichment was also performed on up- or downregulated proteins that belonged to the sepsis-independent effects category (**Figure S6a & S6b**). These proteins matched to several mitochondrial functions, such as metabolism, mitophagy, signalling and protein import, and calcium signalling, but was most pronounced in OXPHOS and mitochondrial transmembrane transport pathways involved with ATP synthesis (**Fig 6B-C**). The DAPs associated with transmembrane transport appear in the small molecule transport group in **Fig 6C**. The heart in particular is enriched in OXPHOS components to due to its high energy demand (Doll et al., 2017). In total, 42 of the known 192 OXPHOS component proteins were altered by the interventions in sepsis (**SFig6_reg-pathways_mitocharta**). In the heart, the sepsis itself did not significantly change the abundance level of the central OXPHOS core subunits of complex I-V (**Fig 6D**). In contrast, all interventions including Mem reduced the levels of the proteins linked to these complexes by an average of 50% in the heart (**Fig 6E**). In leukocytes by contrast, there was an average fold change increase of OXPHOS complexes although the changes were not uniform. In liver, the effects were treatment specific and early administration of Gcc increased the relative amount of all complexes whereas combination treatments lowered the concentration. We also observed a reduction of DAPs associated with cristae that house the OXPHOS components, transmembrane components that are associated with the shuttling of ATP between the mitochondrial matrix and the cytosol, calcium homeostasis and mitophagy. In all cases, GccMem8h had the least sepsis-independent effects (**Fig 6F**). Our data demonstrate an organotypic pattern of mitochondrial dysfunction by interventions. Taken together, these results demonstrate that GccMem8h introduced the least number of mitochondrial pathway impairments in the heart.

Cardiac dysfunction in sepsis has been proposed to be a silent process without major cardiomyocyte apoptosis and is thought to arise due to mitochondrial dysfunction. However, no clear signs of mitochondrial damage were observed except hydropic mitochondria and perturbed cristae. (Takasu et al., 2013). Our analyses reveal that Gcc was largely beneficial in the heart. However, sepsis-independent effects pertaining to ATP generation and calcium ion transport were still present. In contrast, Mem alone or with early administration of Gcc introduced additional sepsis-independent effects including reduction of β-oxidation, cell redox homeostasis, and metabolism of amino acid and derivatives, all of which are involved in ATP generation (Fillmore et al., 2014). In conclusion, our data suggest that interventions could introduce subtle yet potent changes in the mitochondrial proteome in sepsis, which in turn could influence cardiac function. Thus, sepsis-independent effects DAPs represent a previously unexplored and important category that should be considered to score treatment benefits and probe unexpected drug synergies.

## Discussion

Using a combinatorial strategy of conventional markers, histology, proteomics, and systems biology approach, we demonstrate several novel elements of the host response during the course of sepsis and key therapeutic interventions. This approach is more comprehensive than traditional physiological assessments, which are limited to fewer grosser parameters like immunological responses, weight loss, or survival. Previous proteomic strategies rely on a ‘triangular’ approach, where a few targets out of thousands of proteins are chosen to describe pathophysiology and offer the advantage of translation to larger cohorts with high sensitivity. However, in sepsis this approach may be limited in defining disease and intervention complexities due to its reliance on fewer targets. In contrast, we utilized a ‘rectangular’ approach, where maximal proteome coverage allowed for discovery of proteostates and scoring of intervention effects, enabling better insight into underlying biology (Geyer et al., 2017). Recently, the concept of a treatable trait was proposed. We propose that our combinatorial approach presented will aid in more robust stratification and characterization of deeper fine-tuned systemic changes in sepsis that are supported by biologically relevant benchmarked parameters.

Although the plasma is a source of major interest for organ damage biomarkers in sepsis, events regulating their appearance in plasma is largely undefined. We provide 275 tissue-specific candidate proteins that distinguish between early inflammation and late organ damage. These data may strengthen diagnosis in combination with other inflammatory and organ damage markers, and perhaps help distinguish between sepsis and other conditions. Many tissue-enriched proteins appear in plasma at 12 h and 18 h and many are liver-derived. This resembles the sepsis δ phenotype described by Seymour et al. (Seymour et al., 2019), which is characterized by liver failure and elevated IL-6, BUN, and ALT. The liver has a crucial role in regulating both immune and metabolic responses in sepsis (Strnad et al., 2017) and its role in regulating crosstalk in sepsis remains understudied. Apart from being the major contributor to the plasma proteome, the liver plays a key in detoxification, modifying insulin sensitivity, secreting soluble signalling molecules and metabolites, and glucocorticoid, amino acid, lipid, and glucose metabolism (Yan et al., 2014). Disruption of normal liver function can result in conditions like lipotoxicity that affects kidney and heart (Van Wyngene et al., 2020), and accumulation of excess amino acids that can cause brain damage (Felipo and Butterworth, 2002). Our findings have implications for improved characterization of unique host responses over time in organs and the blood compartment and their contribution to organ dysfunction.

Beyond defining organ dysfunction signatures in sepsis, we further used computational strategies to study treatment effects in preventing and reverting this organ damage. We chose the glucocorticoid methylprednisolone and the beta-lactam antibiotic meropenem due to their broad anti-inflammatory and antimicrobial effects, respectively. The combination of glucocorticoids and antibiotics administered at 8 h was most beneficial on account of high number of reversions and minimal side effects. Exogenous glucocorticoid therapy downregulates GR in the liver, and increases levels of lipids in plasma, all of which are detrimental (Jenniskens et al., 2018; Vandewalle and Libert, 2020). Drugs enhancing lipid clearance like PCSK9 inhibitors (Walley et al., 2014) can be envisioned as an adjuvant treatment for glucocorticoids to facilitate lipid clearance in sepsis.

Although, glucocorticoids do not seem to affect overall mortality in the ICU, they were effective in patients with severe immunosuppression (Antcliffe et al., 2019). In our model mortality was not assessed but interventions reduce the heterogeneity among different individuals on the compartments that perform an immune function (**Figure S6d**), whereas organs that do not perform an immune function and are mainly involved in metabolism are heterogeneous. Thus, the interventions exert a greater effect on responder organs in contrast to bystander organs. Additional experiments are required to validate the use of adjuvant drugs to resuscitate metabolic deregulation and responses need to be benchmarked to study their relevance in mortality. Nonetheless, our approach will enable exploration of the link between timing, dose, drug synergies, and corresponding proteotypes in sepsis.

Cardiac dysfunction is an important component in sepsis multiorgan failure and corresponding mitochondrial dysfunction is linked to more severe outcomes (Takasu et al., 2013). However, the role of treatments in preserving or exacerbating cardiac mitochondrial dysfunction is unclear. Our data show that hepatic mitochondrial ATP components including the OXPHOS are affected mainly in the liver but not in the heart. In contrast, antibiotics and glucocorticoids introduced cardiac mitochondrial fluctuations in septic animals only. The role of unintended mitochondrial dysfunction in post-sepsis sequelae, the potential use of mitochondrial resuscitating adjuvants to counter organ dysfunction, and identification of surrogate marker that delineates cardiac mitochondrial status, need to be studied in relevant model systems and patients. Our strategy can be used to probe drug side effects in other model systems.

The major limitation of our study was that our study was performed in mice where only parts of the response are similar to humans and therefore captured. Further validation studies need to be performed with human material. We chose the human isolate *E. coli* O18:K1 as the infecting strain and administered a low dose of 10^4^ bacteria intraperitoneally as described previously (van ‘t Veer et al., 2011). This strategy of bacterial inoculation allows the bacteria to gradually grow to a high number, producing observable multi-organ damage and a wide spectrum of host responses. At a lower challenge dose of 10^3^ CFU, the *E. coli* were rapidly cleared by the animals (data not shown). We saw IFN-β regulatory type-1 interferon responses, fatty acid metabolism, insulin signalling and mitochondrial dysfunction that were benchmarked using accepted pathophysiological parameters including cytokines, circulating immune cell profiles, and organ damage markers, which have all been described in human sepsis (Cavaillon et al., 2020; Huys et al., 2009; Van Wyngene et al., 2018). Data mining using artificial intelligence of large-scale repositories containing patient and mice proteomes should be analysed to begin to address the translational gap.

This study presents a resource containing proteomic atlas cataloguing deep time-resolved proteome maps and treatment scoring strategy in sepsis. We defined several organotypic signatures of inflammatory and metabolic dysregulation, organ crosstalk and damage in sepsis. Further, we characterize treatment effects and provide a scoring strategy that captures reversions, non-reversions, and unintended drug side effects. Thus, improving our understanding of the molecular pathogenesis of organ damage in sepsis and molecular treatment effects on the organism. Finally, we present a novel framework that may be extended to study other diseases, pathogens, treatments, and host response gene knockout functions.

## Supplementary Figures

Figure S1a: Outline of spectral library generation.

Figure S2: Metadata for time course experiments.

Figure S3a: Heatmaps depicting all DAPs observed in organs, plasma and leukocyte during the time course of sepsis.

Figure S3b: Regulation of DAPs in organs, plasma and leukocytes

Figure S3c: Circosplots of GO terms and regulated DAPs in organs, leukocytes and plasma.

Figure S3d: Manually curated heatmap of metascape terms depiciting up- and down-regulated terms during the time course.

Figure S4a: Bacterial load 2h and 8h post-inoculation of bacteria.

Figure S4b: Metadata of the treatment cohort.

Figure S4c: Heatmaps depicting the aggregated up- and down-regulated metascape GO terms in different interventions.

Figure S4d: Quantification of intervention impact based on the slope and R2 values.

Figure S4e: Intervention impact as assessed by R2 values and slope in an independent cohort.

Figure S5a: Regulated Metascape GO terms of drug response networks.

Figure S5b: Heatmap depicting metascape GO terms of the 275 tissue enriched leakage proteins in plasma.

Figure S6a: Downregulated sepsis-independent effects metascape GO terms.

Figure S6b: Upregulated sepsis-independent effects metascape GO terms.

Figure S6c: Mitocarta enrichment terms in the down regulated sepsis-independent effects (SIE) group.

Figure S6d: Highlighting intervention group clusters using UMAPs.

## Materials and methods

### Bacteria and culture conditions

*Escherichia coli* O18:K1 (van ‘t Veer et al., 2011) from a freezer stock (15% glycerol) was grown in Luria-Bertani broth overnight (o/n) at 37°C and 5% CO_2_ without shaking.

### *E. coli* infection in mice

All animal use and procedures were approved by the local Malmö/Lund Institutional Animal Care and Use Committee, ethical permit number 03681-2019. *E. coli* O18:K1 was grown to an optical density of 0.25 at 620 nm in pre-warmed Luria broth (37°C, 5% CO_2_). Bacteria were washed and resuspended in sterile DPBS to 5 × 10^4^ CFU/ml. Nine-week-old female and male C57BL/6J mice (Janvier, Le Genest-Saint-Isle, France) were infected with 200 µl (10^4^ CFU) bacteria by intraperitoneal injection. The control group was similarly injected with sterile DPBS. Body weight and general symptoms of infection were monitored regularly. Mice were sacrificed at 0, 6, 12, and 18 hours post infection (h.p.i.) for the time-course, 2 and 8 hours for the determination of treatment window and finally all animals were sacrificed at 18 hours for treatment with antibiotics and corticosteroids. Blood and organs (liver, lung, spleen, heart and kidney) were collected. Blood was taken by cardiac puncture and collected in tubes containing sodium citrate (MiniCollect tube, Greiner Bio-One). Methylprednisolone (Solu-Medrol™, Pfizer) and meropenem (Pfizer) were injected intravenously at 30 mg/kg and 10 mg/kg, respectively, at 2 or 8 h.p.i.. For interventions were validated in 2 independent cohorts (191210_treatment cohort, n= 44, n=6/treatment group; 200319_validation cohort, n=30, n=6/treatment group).

### Organ preparations

Citrated blood collected from infected and control mice was centrifuged (2000 rcf, 10 min) to obtain platelet-free plasma and the cell pellet containing circulating leukocytes was stored for further processing as described later. Plasma was aliquoted and stored at −80°C. Collected organs were homogenized (MagnaLyzer, Roche) in DPBS using sterile silica beads (1 mm diameter, Techtum). The homogenates were plated for viable counts and saved for proteomics. In some experiments, intact organs were taken for histology (see below). Bacterial load in organs was determined by serial diluting and plating (heart, lung, liver, kidney and spleen homogenates) onto blood agar plates. Colony forming units were counted following o/n incubation (37°C, 5% CO_2_), and are presented as CFU/g of tissue. Remaining organ homogenates were centrifuged (20000 rcf, 10 min, 4°C) and supernatants were immediately transferred and aliquoted into new cryovials. All samples were stored at −80 °C until further analysis. Protein concentration was determined using standard BCA assay (Thermo Scientific) according to manufacturer’s instructions.

### Flow cytometry of blood cells

Citrated blood collected from infected and control mice was diluted with HEPES buffer containing mouse Fc-block (BD Pharmingen). Platelets, monocytes and neutrophils were identified using, anti-CD45 FITC (BD Pharmingen), anti-CD41 FITC (BD Pharmingen), anti-Ly6G PE (BD Pharmingen), anti-Ly6C APC-R700 (BD Horizon) and anti-CD11b PerCP (BD Pharmingen). All antibodies were diluted 1:200 and incubated for 15 min at room temperature). Samples were lysed using 1-step Fix/Lyse Solution (e-Bioscience), washed (500 rcf, 5 min), and cell pellets were resuspended in PBS. The samples were analyzed using an Accuri C6 Plus flow cytometer (BD Biosciences), and the data was analyzed using C6 Plus Software (BD Biosciences). Total leukocytes were gated according to characteristic forward and side scatter. Neutrophils were identified as CD11b high, Ly6G high and Lyg6C low. Monocytes were identified as CD11b high, Ly6G negative and Lyg6C intermediate/high.

### Cytokines and organ damage markers

Cytokines in citrated plasma samples were quantified using the BD Cytometric Bead Array Mouse inflammation kit (552364, BD Biosciences) according to manufacturer’s instructions and analyzed using a FACSVerse flow cytometer (BD Biosciences). Plasma levels of alanine aminotransferase (ALT), lactate dehydrogenase (LDH), blood urea nitrogen (BUN), and were measured using ALT Activity Assay Kit (ab105134, Abcam), LDH Assay Kit (ab102526, Abcam) BUN Colorimetric Detection Kit (EIABUN, Invitrogen, Thermo Fisher Scientific), and cardiac troponin I (Abcam, ab285235) respectively. In organ homogenates, levels of IL-6 (88-7064), IL-10 (88-7105), TNF-α (88-7324), IFN-γ (88-7314), and IFN-α (BMS6027) were measured using mouse ELISA kits (Invitrogen, Thermo Fisher Scientific). All ELISA and colorimetric assays were performed according to manufacturer’s instructions in combination with a microplate reader (VICTOR3 Multilabel Plate Reader, Perkin Elmer).

### Pathology scoring and immunohistochemistry of organs

Heart, lung, liver, spleen, and kidney tissue samples from infected and control mice were fixed in Histofix (Histolab Products AB, Askim, Sweden) for 48 h, dehydrated in 70% EtOH for at least 24 h, then embedded into paraffin blocks and sectioned (4 μM microtome, Leica RM2255). Sections were transferred to slides and deparaffinized (60°C, 30 min) followed by dehydration in Histolab Clear (#14250, Histolab). Sections were stained using Mayer’s hematoxylin and eosin (H&E) (Histolab Products AB, Askim, Sweden). Sections were air-dried at room temperature and mounted with Pertex 5 (#00840, Histolab) and a glass coverslip (1.5 mm). Imaging was performed using a light microscope (Nikon Eclipse 80i: 10x magnification in skin and 40x magnification in kidney and liver). Representative images from H&E-stained sections of liver, spleen and kidneys were acquired at 20x and 40x magnifications. Image analysis of the sections was performed with FIJI version 1.53. For quantification of liver dysfunction, the presence of clots in the liver vessels was counted manually. For quantification of kidney dysfunction, mesangial proliferation in kidneys was assessed. A minimum of 36 glomeruli were selected using the freehand tool and processed similarly as described for spleen above, with Huang chosen as the method for auto threshold. Images were deconvolved using the color deconvolution function, with vectors set to H&E. The image depicting hematoxylin staining was selected and auto threshold set to Otsu was applied. The thresholded image was added to the ROI manager and then objects within the ROI were measured.

### Leukocyte preparation

The blood cell pellet obtained post collection of plasma was transferred to a fresh 15ml Falcon tube (Sarstedt, Germany) and subjected to erythrocyte lysis using buffer EL (Qiagen, 5ml) for 10 minutes on ice, followed by centrifugation @500g for 10 minutes (4 ° C, acc/dec = 9). The supernatant was collected carefuly without disturbing the pellet and discarded. The process was repeated twice (3 in total) and the leukocyte enriched pellet was lysed in 350 µl RLT buffer (Qiagen) and stored at −20 °C. Cell lysates were processed using the Allprep DNA/RNA kit (Qiagen). The protein containing flow-through (500µl) obtained after total RNA isolation was subjected to precipitation suing 3 volumes (1.5ml) of ice-cold acetone for 1 hour in the freezer. Precipitates were then centrifuged for 15 minutes @14000g, 4°C. Supernatant was discarded and the samples were air dried at room temperature for 30 minutes. The protein precipitates were then dissolved in 50 µl 0.1% RapiGest SF (Waters), 8M urea (Sigma Aldrich) and 100mM ammonium bicarbonate, pH=8 (Sigma-Aldrich) and stored at −80°C. 25 µg Protein was takenm for in-solution digest as described below. For the leukocyte data present in the time course experiments were added from an independent cohort.

### MS sample preparation and data acquisition

Plasma, leukocytes and organ homogenates (liver, kidney, spleen, heart and lung) were denatured with 8 M urea and reduced with 5 mM Tris(2-carboxyethyl)phosphine hydrochloride, pH 7.0 for 45 min at 37 °C, and alkylated with 25 mM iodoacetamide (Sigma) for 30 min followed by dilution with 100 mM ammonium bicarbonate to a final urea concentration below 1.5 M. Proteins were digested by incubation with trypsin (1/100, w/w, Sequencing Grade Modified Trypsin, Porcine; Promega) for at least 9 h at 37°C. Digestion was stopped using 5% trifluoracetic acid (Sigma) to pH 2 to 3. Peptide clean-up was performed by C18 reversed-phase spin columns according to manufacturer instructions (Silica C18 300 Å Columns; Harvard Apparatus). Solvents were removed using a vacuum concentrator (Genevac, miVac) and samples were resuspended in 50 μl HPLC-water (Fisher Chemical) with 2% acetonitrile and 0.2% formic acid (Sigma). Peptide analyses (corresponding to 1 µg protein) were performed on a Q Exactive HF-X mass spectrometer (Thermo Fisher Scientific) connected to an EASY-nLC 1200 ultra-HPLC system (Thermo Fisher Scientific). Peptides were trapped on precolumn (PepMap100 C18 3 μm; 75 μm × 2 cm; Thermo Fisher Scientific) and separated on an EASY-Spray column (ES903, column temperature 45 °C; Thermo Fisher Scientific). Equilibrations of columns and sample loading were performed per manufacturer’s guidelines. Mobile phases of solvent A (0.1% formic acid), and solvent B (0.1% formic acid, 80% acetonitrile) was used to run a linear gradient from 5% to 38% over 90 min at a flow rate of 350 nl/min. The variable window data in-dependent acquisition (DIA) method is described by Bruderer et al (Bruderer et al., 2017). The data dependent acquisition (DDA) method was the manufacturer’s default for ‘high sample amount’. LC-MS performance was quality controlled with yeast protein extract digest (Promega). MS raw data was stored and managed by openBIS (v20.10.0) (Bauch et al., 2011) and converted to centroid indexed mzMLs with ThermoRawFileParser (v1.2.1) (Hulstaert et al., 2020).

### Spectra library generation

Mouse peptide samples (plasma, spleen, kidney, liver, leukocytes and heart) from the time course experiment were pooled per sample type and time points (T0 together with T6 and T12 together with T18) and then fractionated with High pH Reversed-Phase (Pierce) into totally 96 samples, Theses samples were spiked with PROCAL standard peptides (Zolg et al., 2017) and were analyzed with DDA, The resulting MS data was searched with FragPipe (v12.2) (Kong et al., 2017; Teo et al., 2021) using default parameters against a protein sequence database containing the mouse reference proteome (EMBL-EBI RELEASE 2019_04), contaminants and PROCAL together with decoys (in total 44824 entries) using MSFragger (v2.4) and Philosopher (v3.2.3) applying 1% FDR. A spectral library from was compiled from the FragPipe output using Spectrast (v5.0), and msproteomicstools (v0.11.0) for retention time (RT) normalization of the PROCAL precursors. The library contained 11,510 proteins, 152,510 precursors, and 152,299 transitions together with corresponding decoys generated with OpenSwathDecoyGenerator v2.4.

### DIA data extraction

A RT normalization library was generated by extracting all albumin precursors from the library generated above and then refined by testing the precursors for data missingness and delta RT scores in a subset of DIA files with OpenSwathWorkflow (v2.4). 59 albumin precursors were finally selected for the final RT normalization library and used for all DIA data extraction downstream. The main searches and scorings were performed by sample type and mouse experiment. DIA data files (n=568) were analyzed against the RT and main spectral libraries with OpenSwathWorkflow (v2.4). The OpenSwath data was scored with PyProphet (v2.1.3) utilizing 1% FDR on protein and peptide levels. Finally, the peakgroups were aligned with TRIC from msproteomicstools (v0.11.0).

### Bioinformatics analysis

MS data analysis was performed using custom scripts in R (3.6) with the R package collection Tidyverse (1.3.0) and Bioconductor package manager BiocManager (v3.9). 17,881,324 rows of TRIC output were read into R and decoy precursors were removed. Precursor intensities were summed into protein intensities and the data was normalized with function justvsn() from package vsn (v3.52.0) per sample type and mouse experiment. Proteins with missing values were imputed with base R stats::runif() function with the assumption of left-censored data missing not at random. Differential abundance testing was performed with R package limma (1.11.1) or with base R function stats::t.test(). Cut-offs were foldchange > ±1.5 and adjusted (with method of Benjamini, Hochberg) p-value < 0.05. Functional and pathway enrichment analysis was performed with Metascape (v33) (Zhou et al., 2019) using the web interface (https://metascape.org/) and the ‘Express Analysis of Multiple Gene Lists’ workflow. Metascape results were downloaded as ‘zip-files’ and protein members from terms together with enrichment results were extracted with custom R scripts. The batch-effect correction approach was to utilize protein log2 foldchange in relation to naïve (h.p.i. = 0) samples per experiment. UMAP projections were generated with R package uwot (v0.1.4). Heatmaps were generated with R package ComplexHeatmap (v2.0.0) using ward.D2 cluster analysis. The MitoCarta3 mouse database was used for defining mitochondrial proteins and subdivision into functional groups (Rath et al., 2021).

### Statistical analysis

Statistical analysis was, unless otherwise stated, performed using nonparametric Mann-Whitney tests (Prism v9.1.0 software; GraphPad, Inc). P values < 0.05 were considered statistically significant.

## Acknowledgements

Gisela Hovold, Louise Thelaus and Jane Fisher are acknowledged for their excellent technical assistance.

