## Supplementary material for "A pharmacoproteomic landscape of organotypic intervention responses in Gram-negative sepsis": Figure_S1

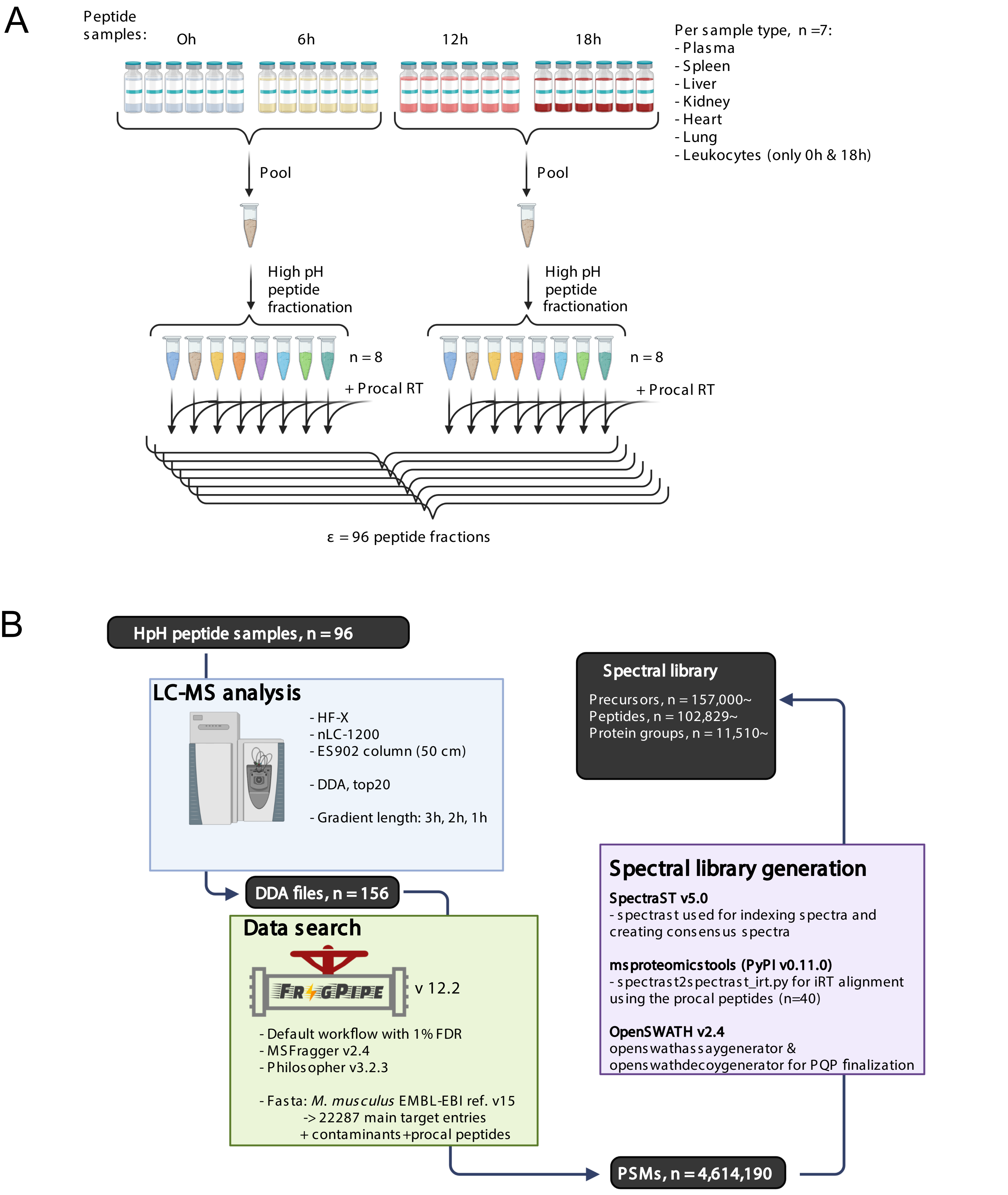


Figure S1 Outline of spectral library generation

A) Organs homogentaes, leukocytes and plasma were pooled and fractionated using high pH peptide fractionation (Pierce), spiked with retention time peptides (Procal RT)

B) LC MS analysis and data seraching and creaion of spectral library creation (See methods for details)
