## Supplementary material for "A pharmacoproteomic landscape of organotypic intervention responses in Gram-negative sepsis": Figure_S2

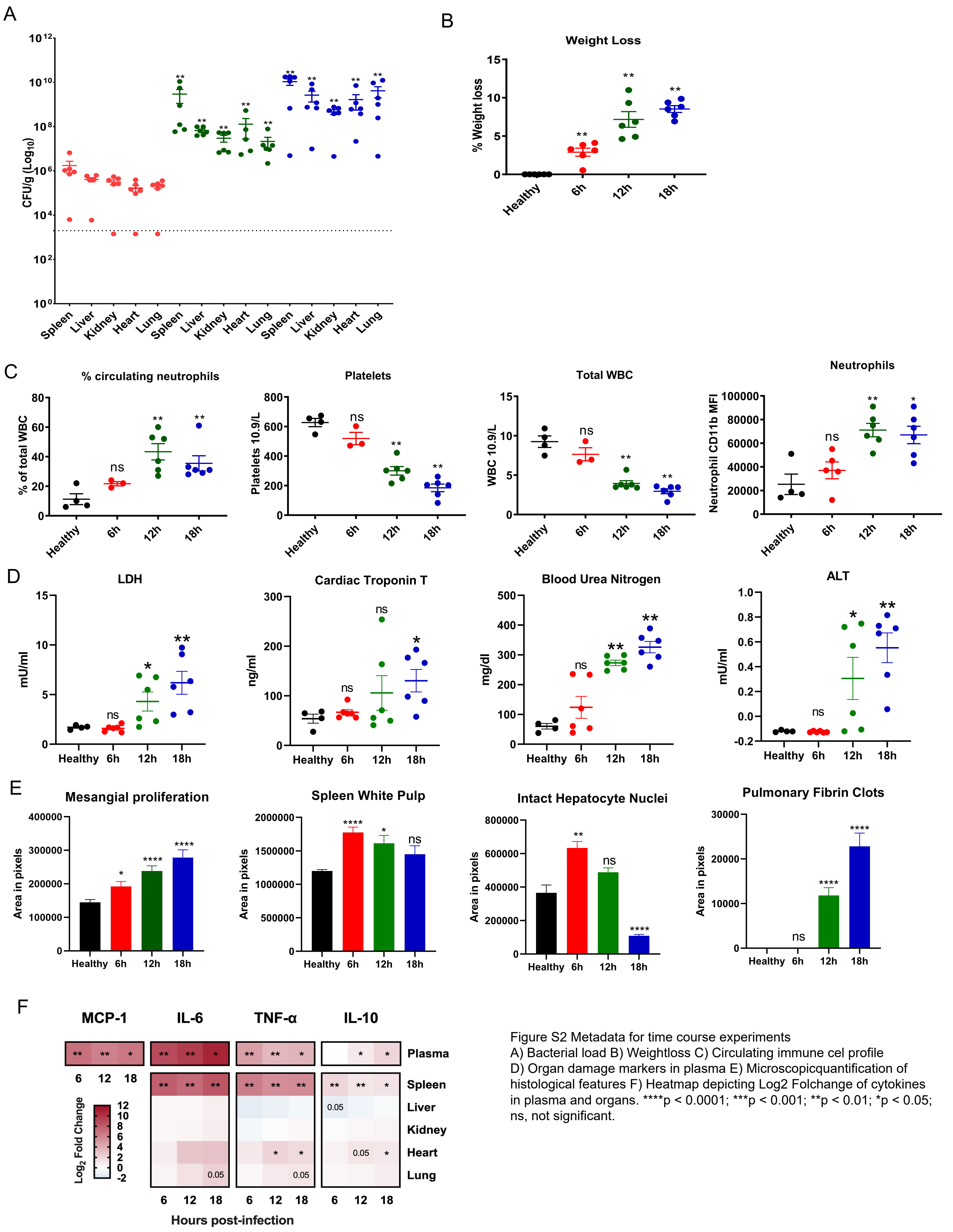


Figure S2 Metadata for time course experiments
A) Bacterial load B) Weightloss C) Circulating immune cel profile
D) Organ damage markers in plasma E) Microscopicquantification of histological features F) Heatmap depicting Log2 Folchange of cytokines in plasma and organs. ****p < 0.0001; ***p < 0.001; **p < 0.01; *p < 0.05; ns, not significant.
