## Supplementary material for "A pharmacoproteomic landscape of organotypic intervention responses in Gram-negative sepsis": Figure_S3a

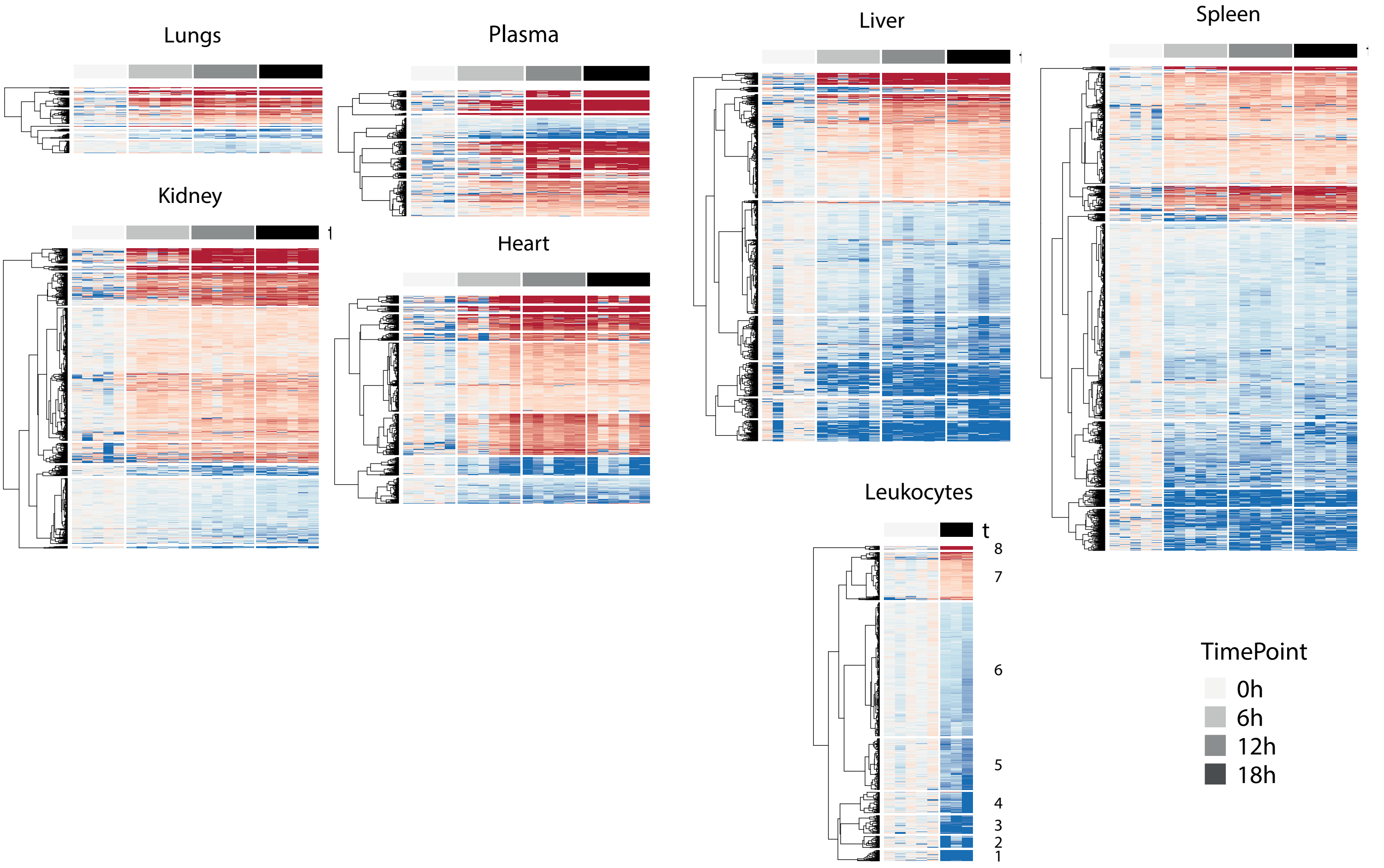


Figure S3a Heatmaps depicting all DAPs observed in organs, plasma and leukocyte during the time course of sepsis.
