## Supplementary material for "A pharmacoproteomic landscape of organotypic intervention responses in Gram-negative sepsis": Figure_S3b

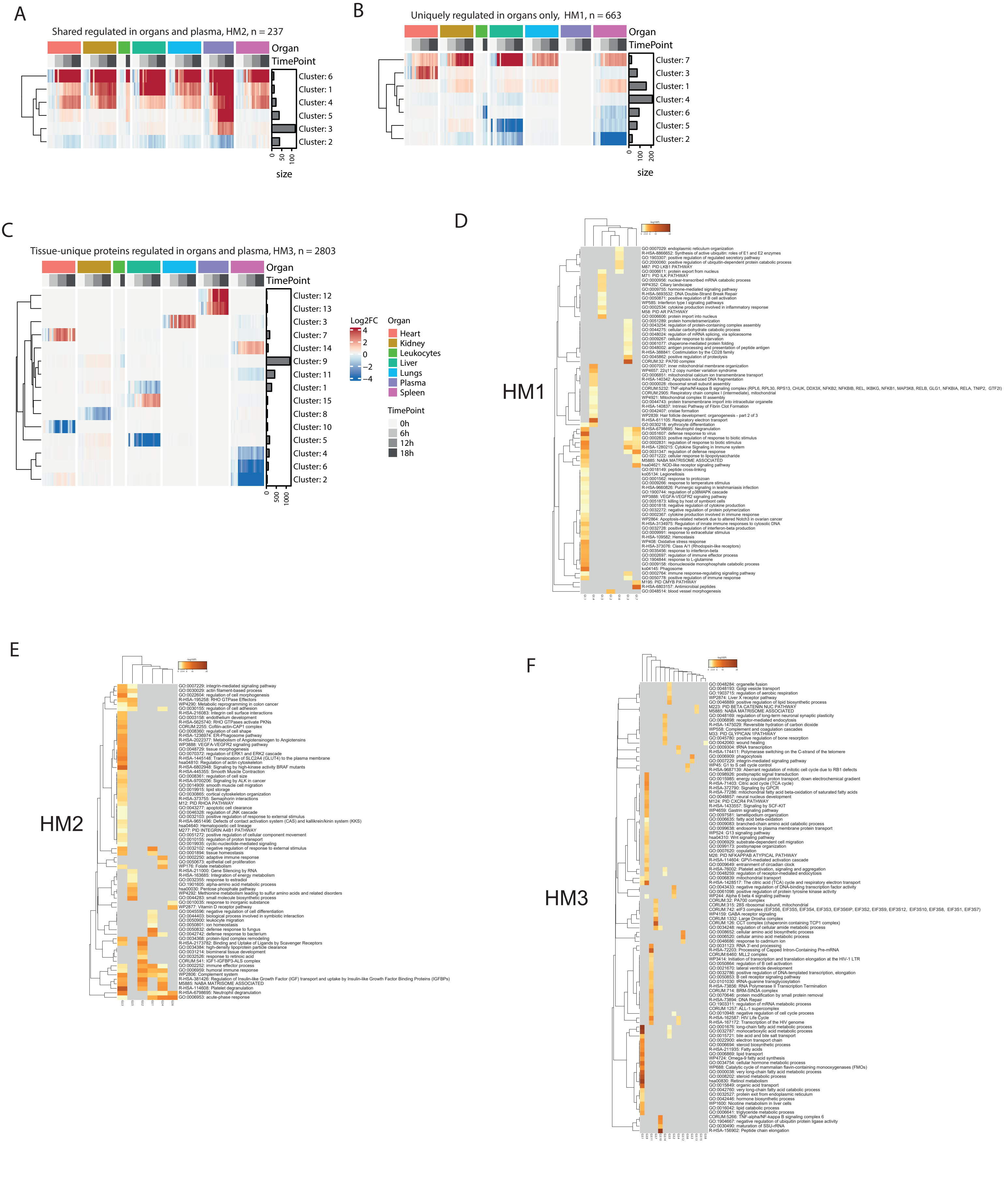


Figure S3b Regulation of DAPs in organs, plasma and leukocytes
A) HM2, Shared DAPs regulated in atleast 1 organ, plasma and leukocytes.
B) HM1, Regulated DAPs shared across organs but not plasma.
C) HM3, Regulated DAPs shared across organs and plasma.
D) Metascape GO terms of cluster HM1, tissue unique DAPs upregulated in organs only.
E) Metascape terms of cluster HM2, shared regulated in organs and plasma
F) Metascape terms for tissue-unique proteins regulated in organs and plasma, HM3.
