## Supplementary figures and images for "A pharmacoproteomic landscape of organotypic intervention responses in Gram-negative sepsis"

### Figure_S3c

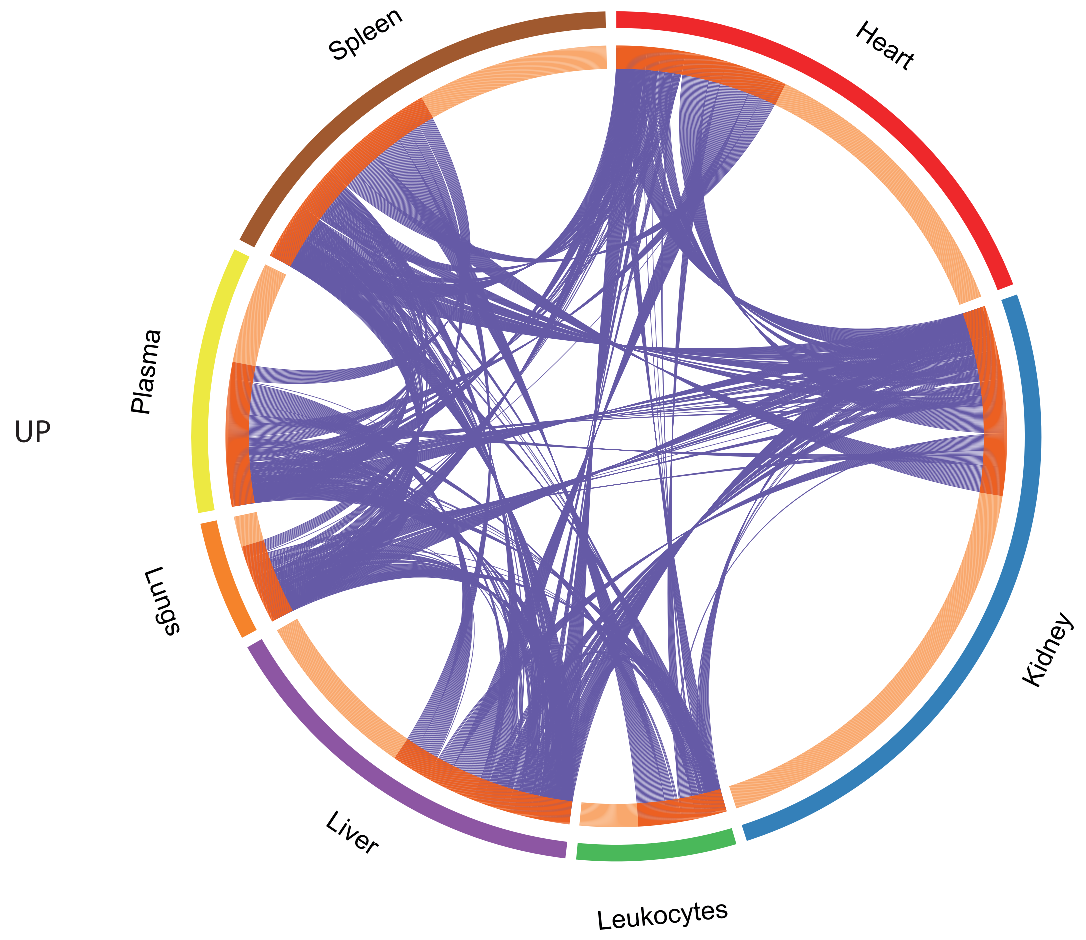


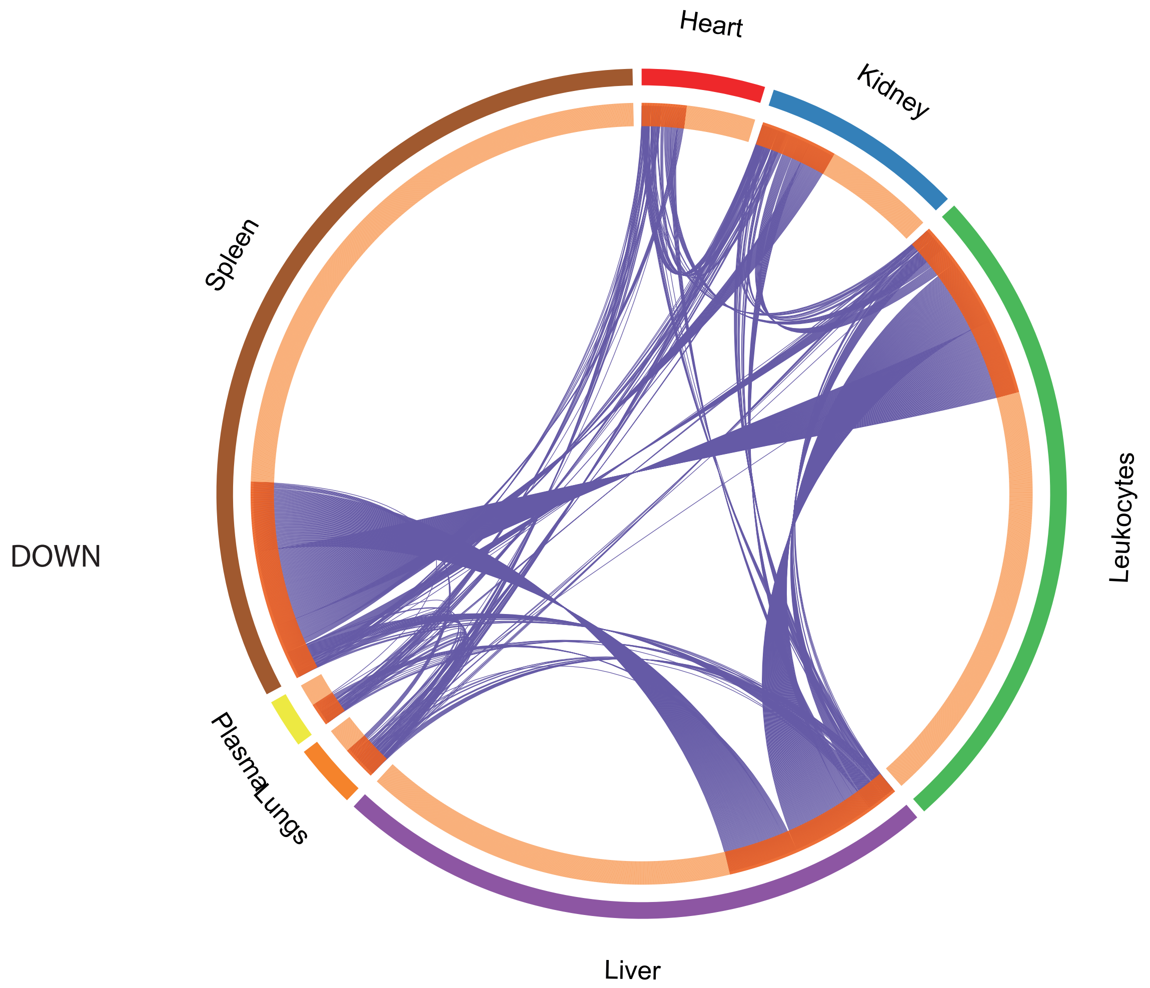


Figure S3c Circosplots of GO terms and regulated DAPs in organs, leukocytes and plasma.

### Figure_S5b

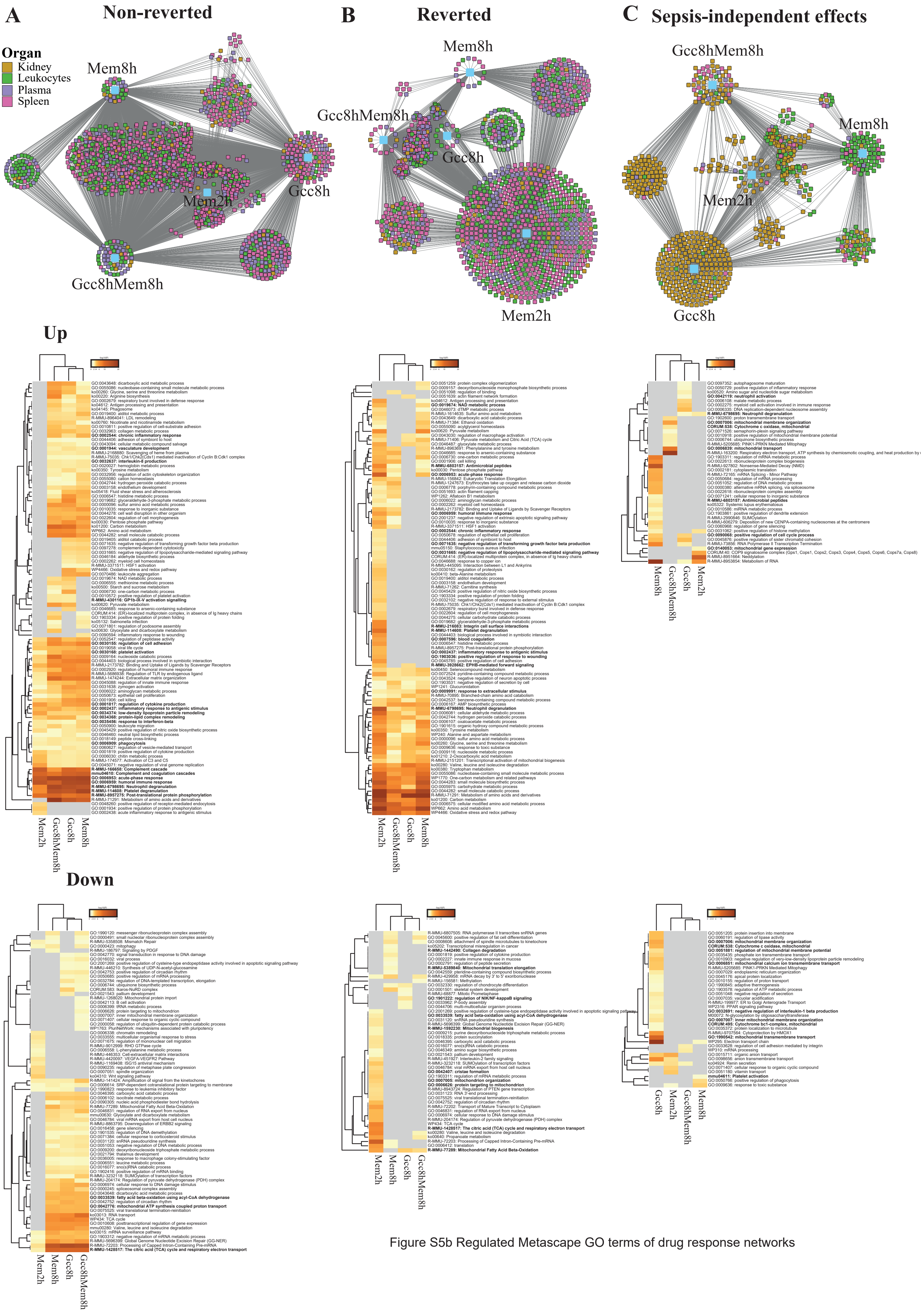


Figure S5b Regulated Metascape GO terms of drug response networks
