## Supplementary material for "A pharmacoproteomic landscape of organotypic intervention responses in Gram-negative sepsis": Figure_S3d

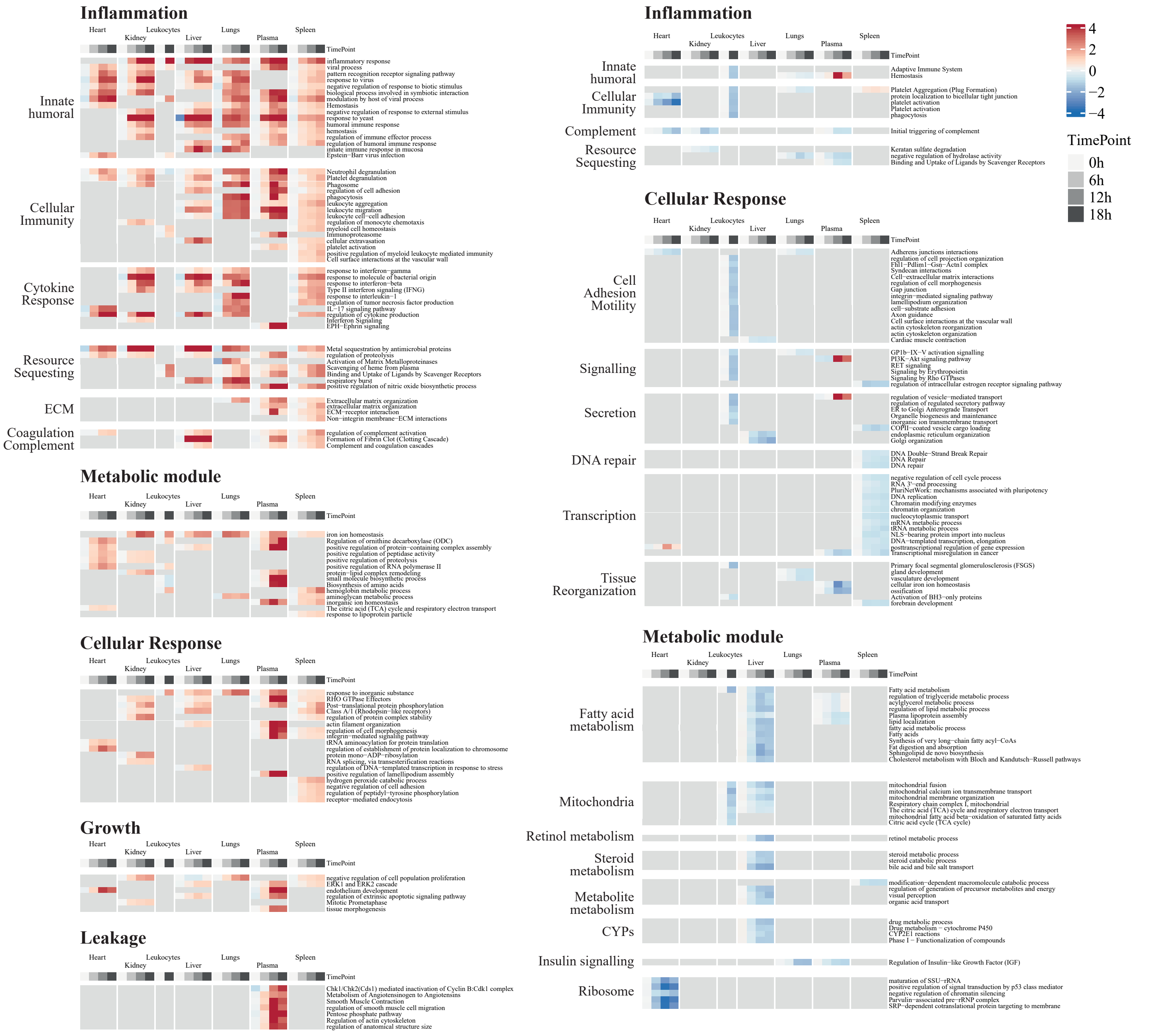


Figure S3d Manually curated heatmap of metascape terms depiciting up- and down-regulated terms during the time course.
