## Supplementary material for "A pharmacoproteomic landscape of organotypic intervention responses in Gram-negative sepsis": Figure_S4a

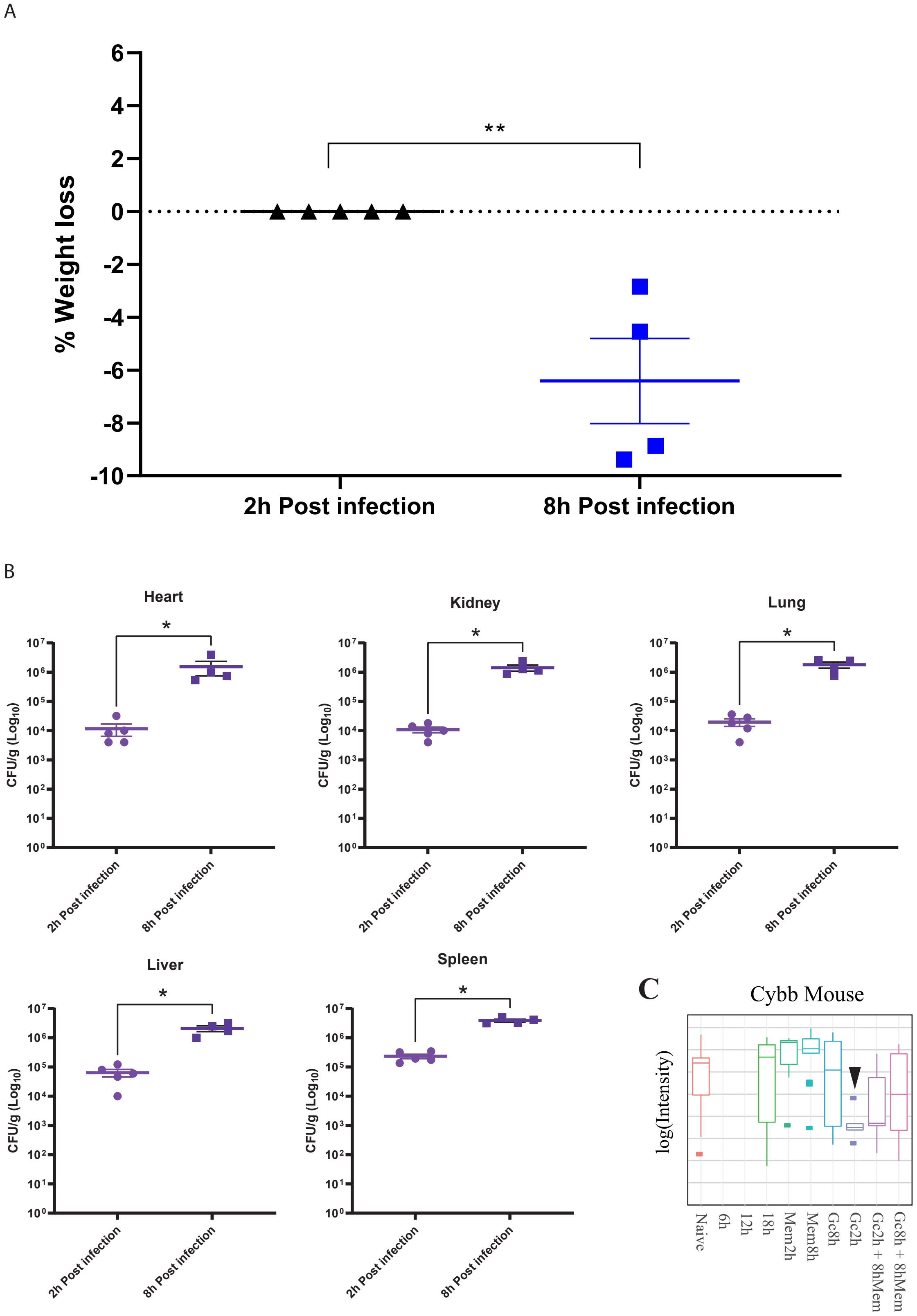


Figure S4a Bacterial load 2h and 8h post-inoculation of bacteria
A) Weight loss B) Bacterial load C) Cybb log intensity levels in leukocytes in the treatment cohort.

****$p < 0.0001$; ***$p < 0.001$; **$p < 0.01$; *$p < 0.05$; ns, not significant.
