## Supplementary material for "A pharmacoproteomic landscape of organotypic intervention responses in Gram-negative sepsis": Figure_S4b

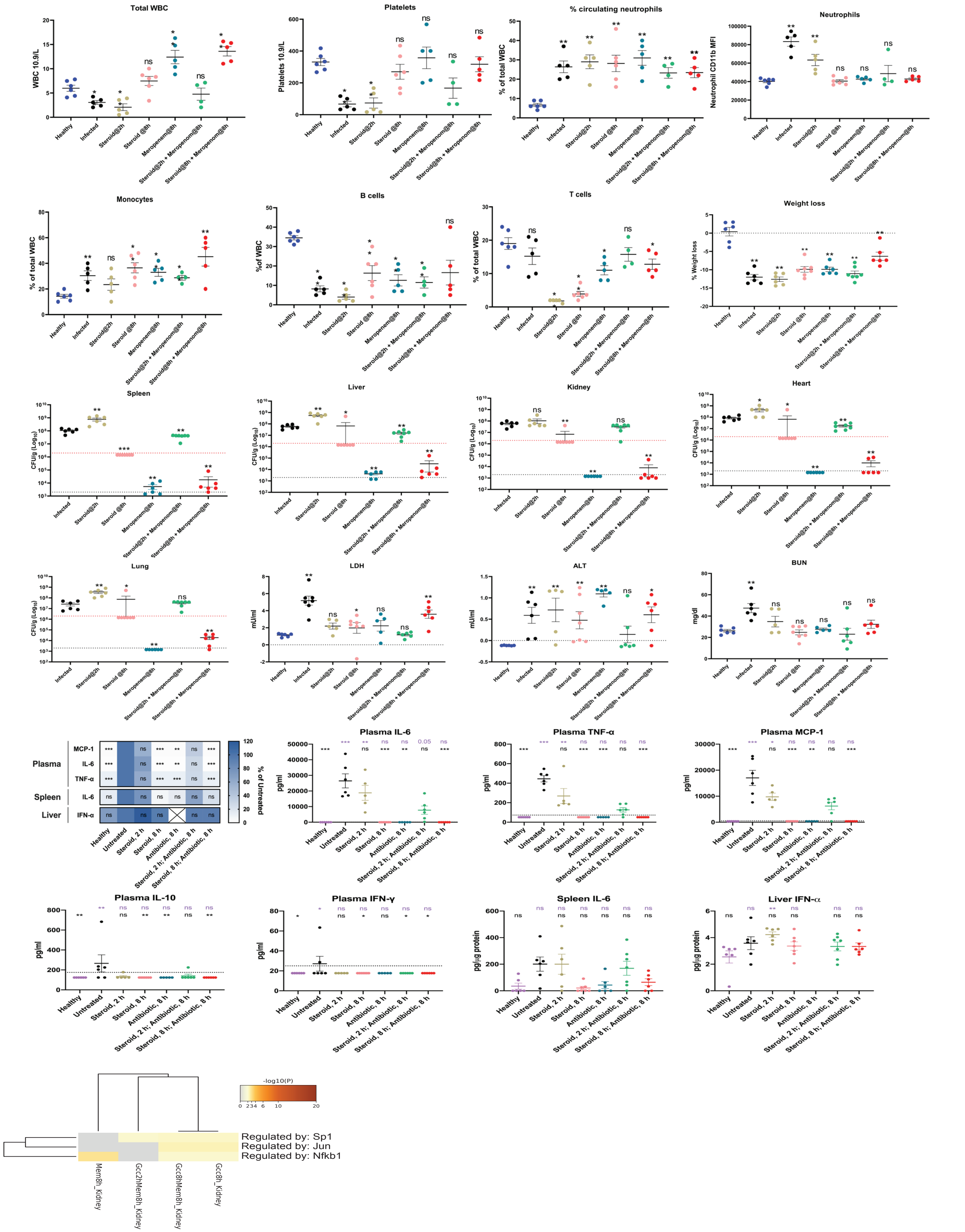


Figure S4b Metadata of the treatment cohort

Physiological parameters of circulating immune cells, organ bacterial load, weightloss, organ damage markers and cytokines in plasma, liver and spleen.

Scatter plots depicting raw levels of cytokines (pg/ul or pg/ug, wherever indicated) used for constructing the heatmap are shown beside the heatmap.

****p < 0.0001; ***p < 0.001; **p < 0.01; *p < 0.05; ns, not significant.

The term steroid was used interchangably with glucocorticoid (Gcc), and antibiotic with meropenem (Mem) in all panels of the figure, except the TRRUST network.

TRRUST networks from metascape analysis showing transcription factors treatment impact of GccMem8h on kidneys
