## Supplementary material for "A pharmacoproteomic landscape of organotypic intervention responses in Gram-negative sepsis": Figure_S4c

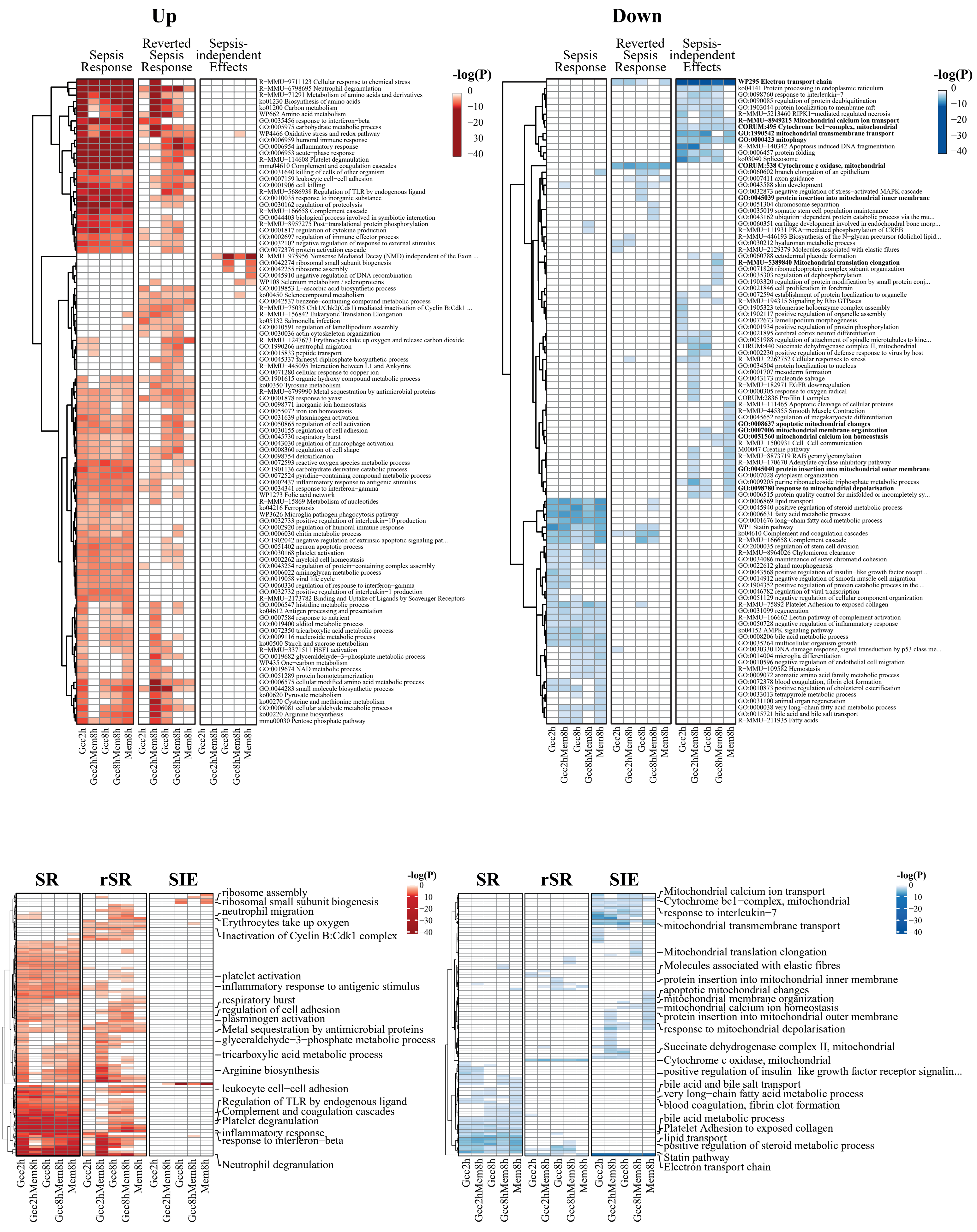


Figure S4c Heatmaps depicting the aggregated up- and down-regulated metascape GO terms in different interventions
SR = Sepsis response, rSR = reverted sepsis response and SIE = Sepsis-independent effects
