## Supplementary material for "A pharmacoproteomic landscape of organotypic intervention responses in Gram-negative sepsis": Figure_S4d

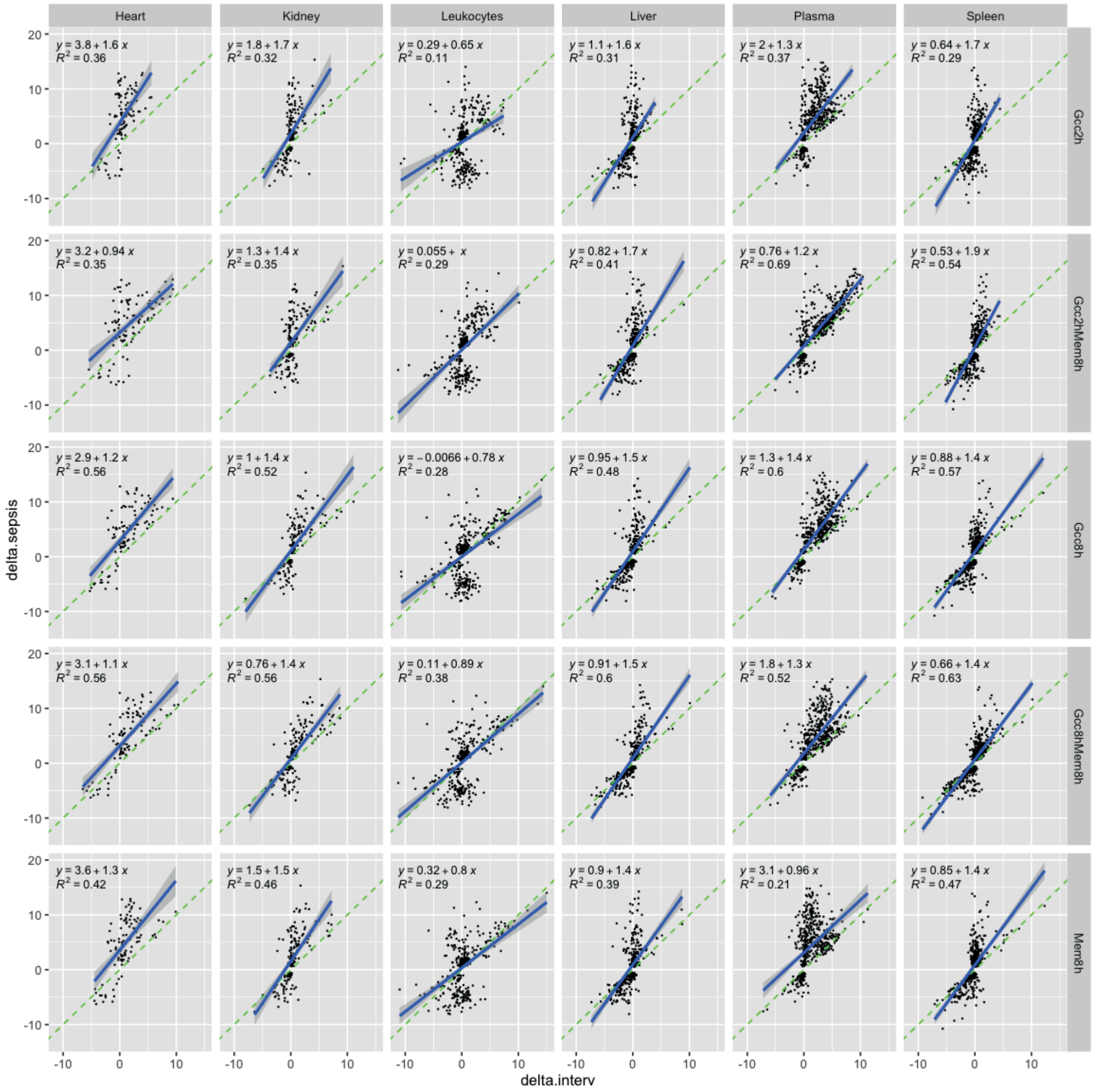


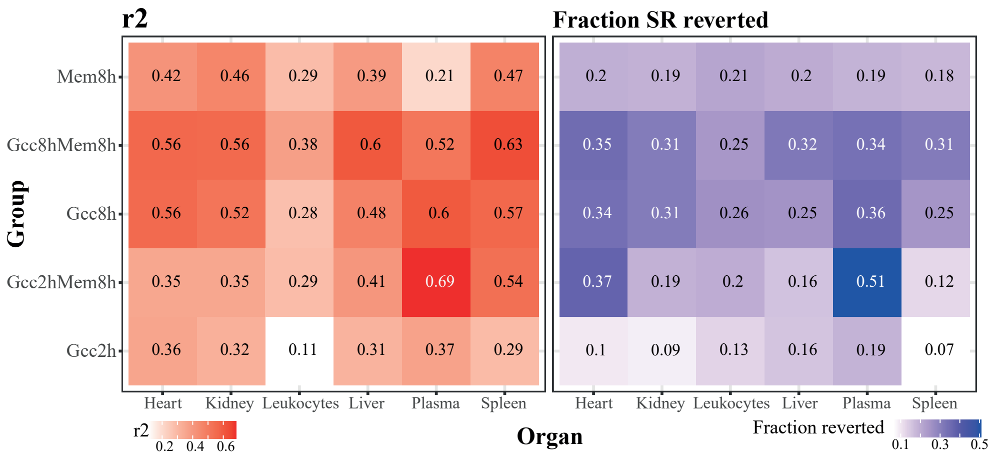


Figure S4d Quantification of intervention impact based on the slope and R2 values
Upper panel depicts lots depicteing the slope and R2 values across organs, plasma and leukocytes.
Lower panel depicts a heat map and values only of R2 values and the fraction reverted (Ratio of Reverted DAPs/Total response)
SR reverted = Sepsis response reverted
