## Supplementary material for "A pharmacoproteomic landscape of organotypic intervention responses in Gram-negative sepsis": Figure_S4e

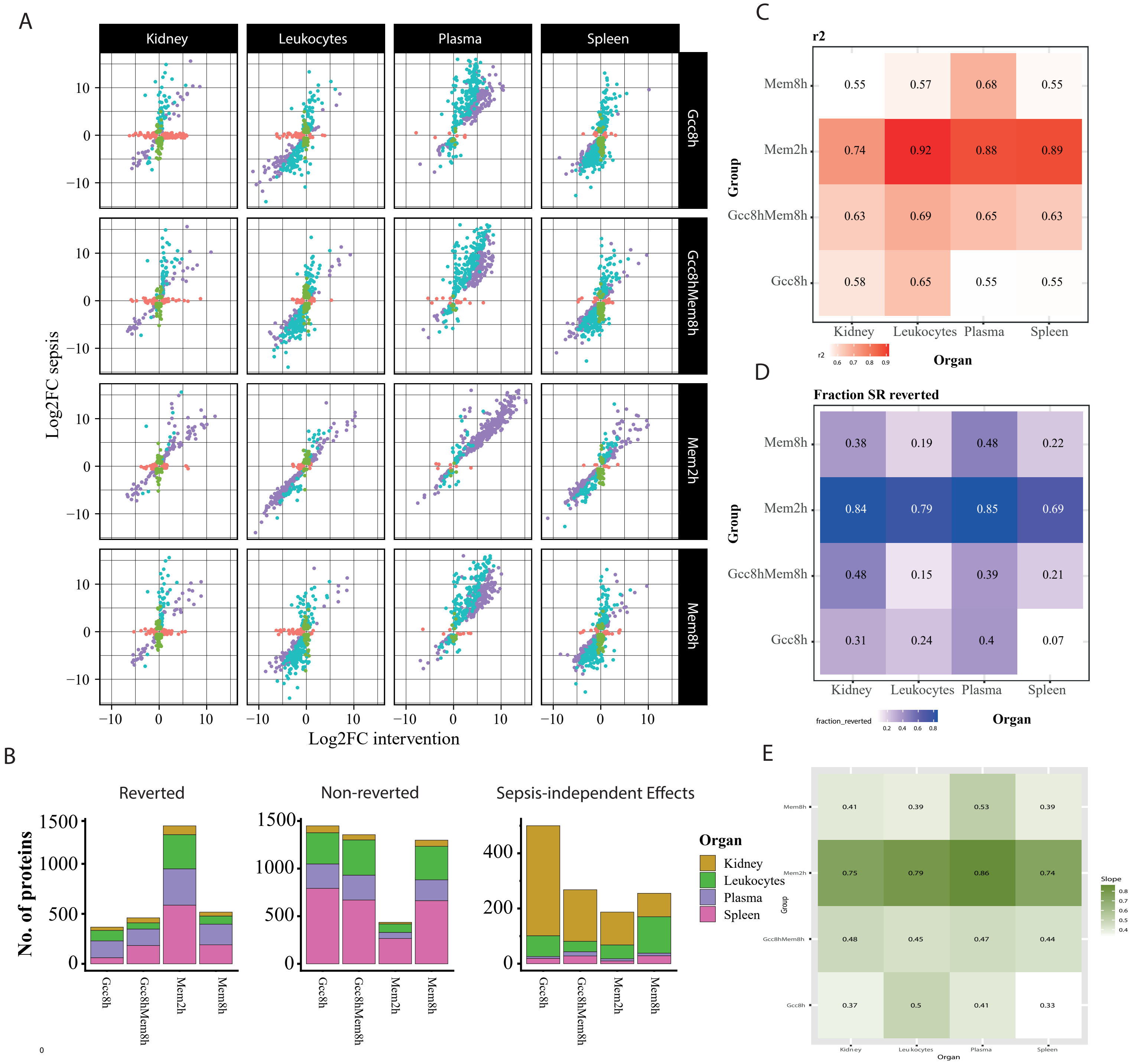


Figure S4e Intervention impact as assessed by R2 values and slope in an independent cohort

A) Plots depiciting reversions, non-reversions and side effects in an independent intervention cohort.

B)Number of proteins in organs across reversions, non-reversions and sepsis-independent effects.

C) R2 values depicted as a heatmap

D) Fraction of reversion by interventions

(Fraction SR reverted = Reversions/total response)

E) Heatmap depiciting values of the slope of reversion

SR reverted = Sepsis response reverted
