## Supplementary material for "A pharmacoproteomic landscape of organotypic intervention responses in Gram-negative sepsis": Figure_S5a

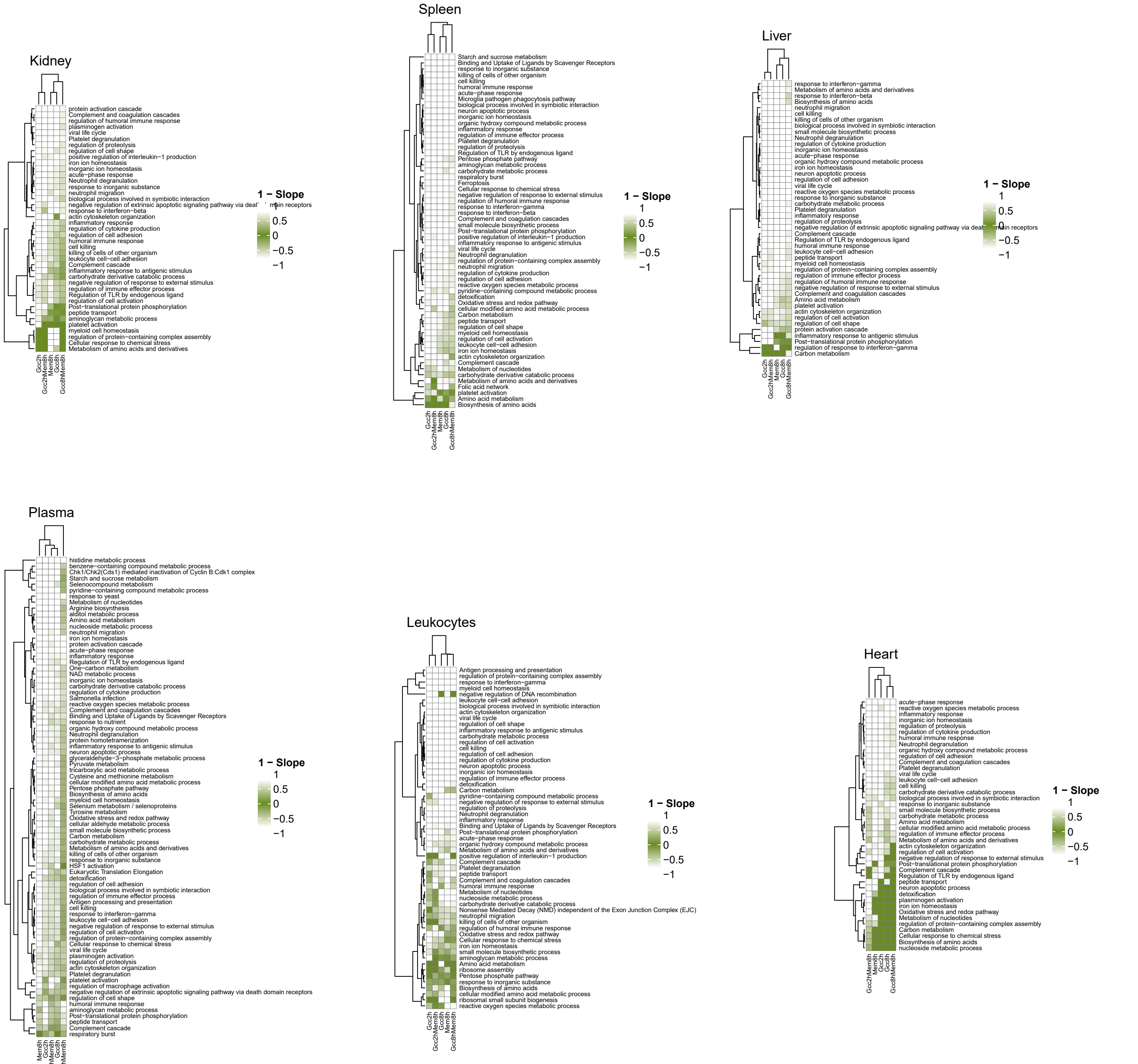


Figure S5a Heatmaps showing 1 - Slope for all interventions across all organs
Heatmaps show the level of reversion of proteins belonging to various metascape GO terms by interventions in all organs. A 1-slope value was calculated for each protein category, where protein categories close to 0 are colored indicating the intervention effect per treatment group shown as a heatmap for functional groups associated with increased protein abundance.
