## Supplementary material for "A pharmacoproteomic landscape of organotypic intervention responses in Gram-negative sepsis": Figure_S5c

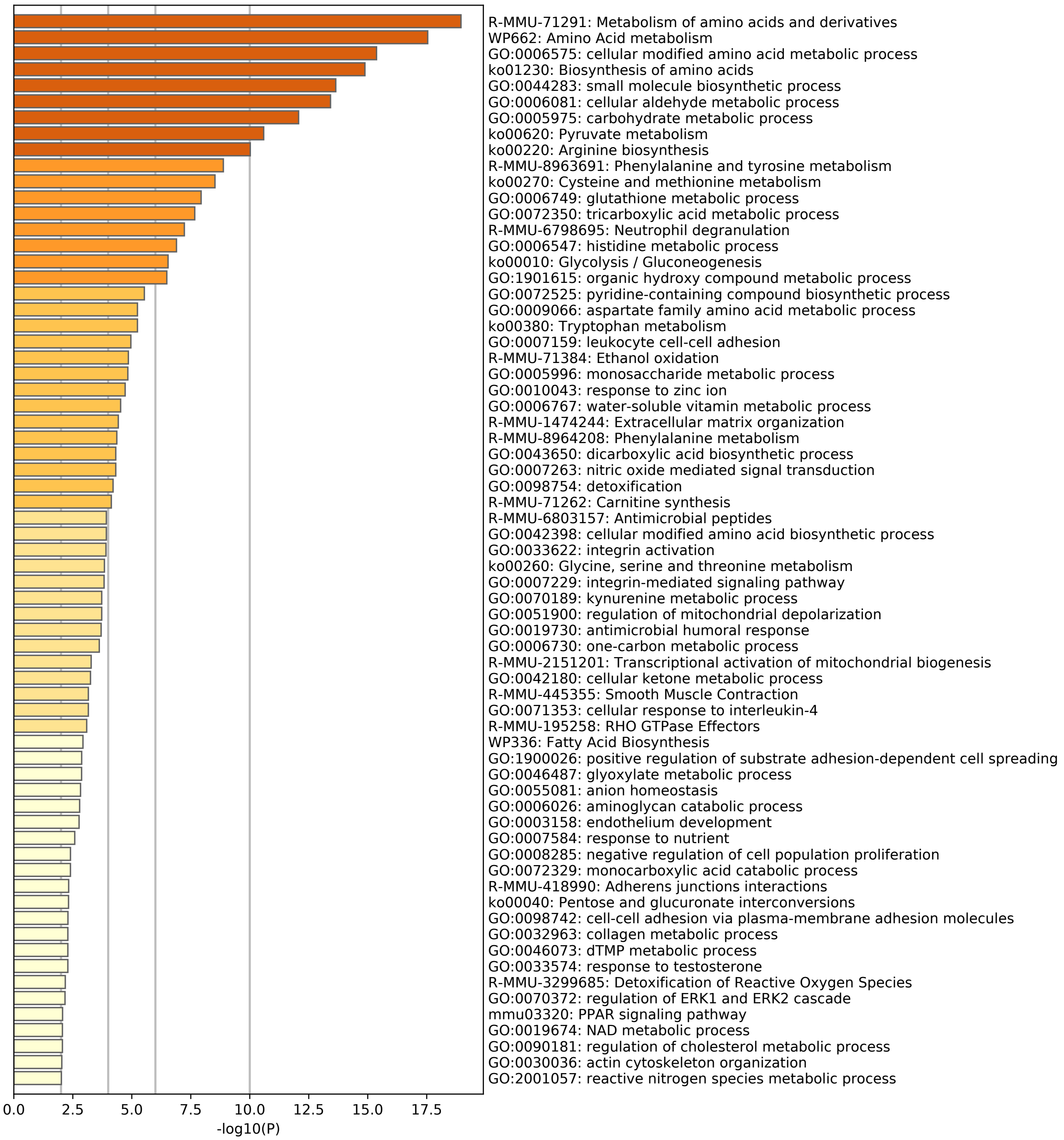


Figure S5c - Heatmap depicting metascape GO terms of the 275 tissue enriched leakage proteins in plasma.
