## Supplementary material for "A pharmacoproteomic landscape of organotypic intervention responses in Gram-negative sepsis": Figure_S6a

Sepsis-independent effects Down

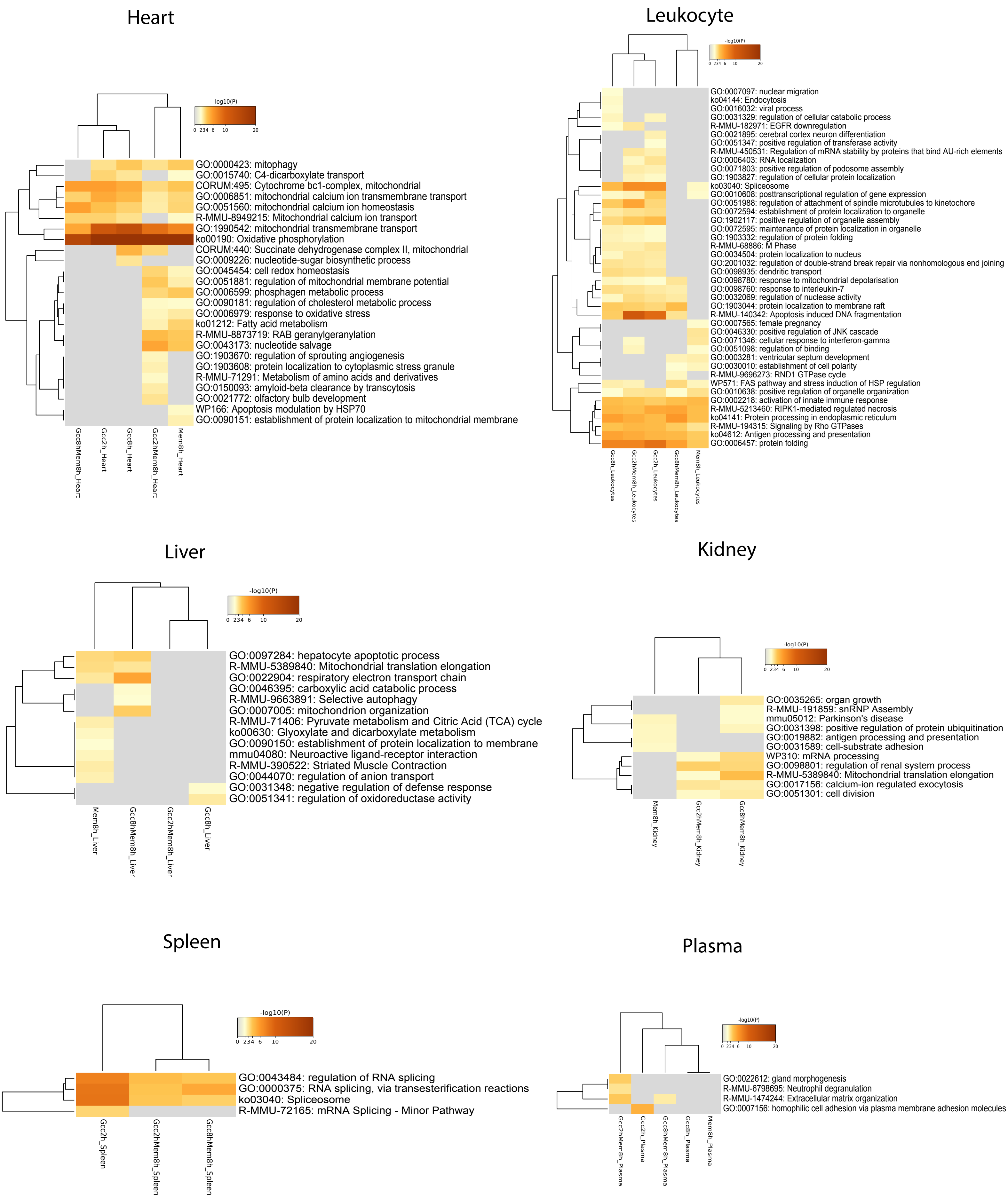


Figure S6a Downregulated sepsis-independent effects metascape GO terms
