## Supplementary material for "A pharmacoproteomic landscape of organotypic intervention responses in Gram-negative sepsis": Figure_S6b

### Sepsis-independent effects Up

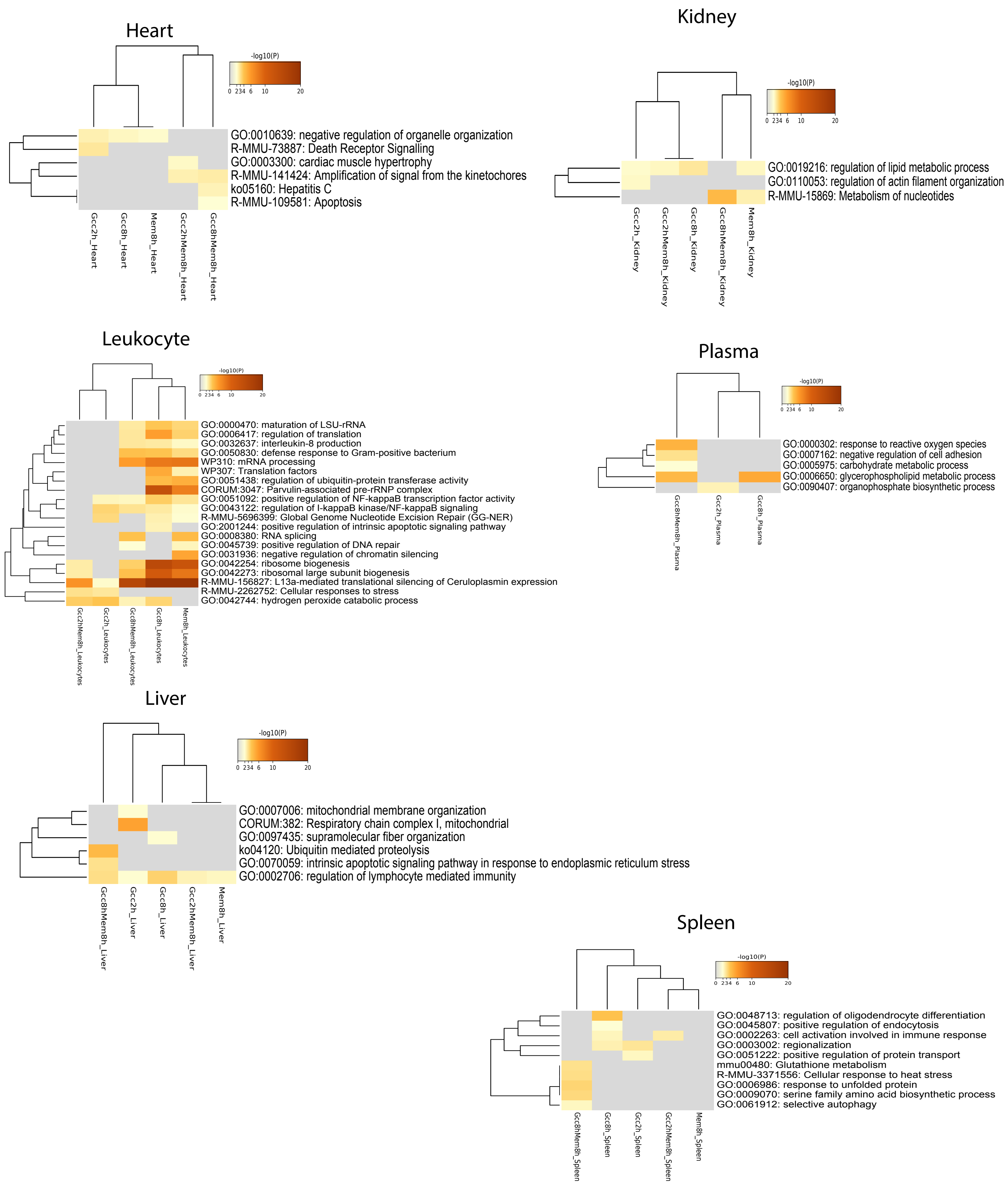


Figure S6b Upregulated sepsis-independent effects metascape GO terms
