## Supplementary material for "A pharmacoproteomic landscape of organotypic intervention responses in Gram-negative sepsis": Figure_S6c

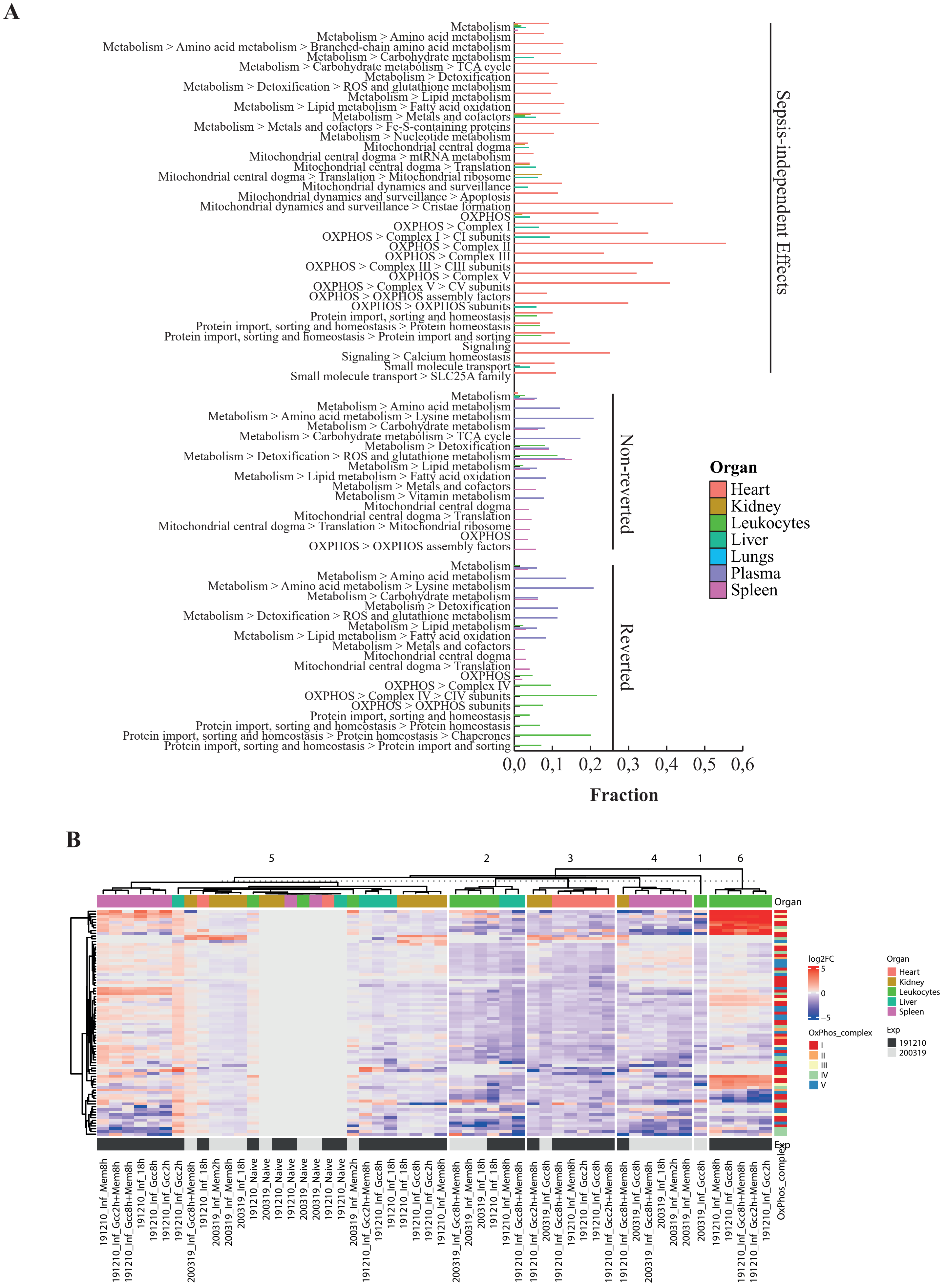


Figure S6c Mitocarta enrichment terms in the down regulated sepsis-independent effects (SIE) group
A) Fraction of the proteins found in mitocarta GO terms in the down regulated SIE group
B) Heatmap depicting log2Fc of OXPHOS component across organs
