## Supplementary material for "A pharmacoproteomic landscape of organotypic intervention responses in Gram-negative sepsis": Figure_S6d

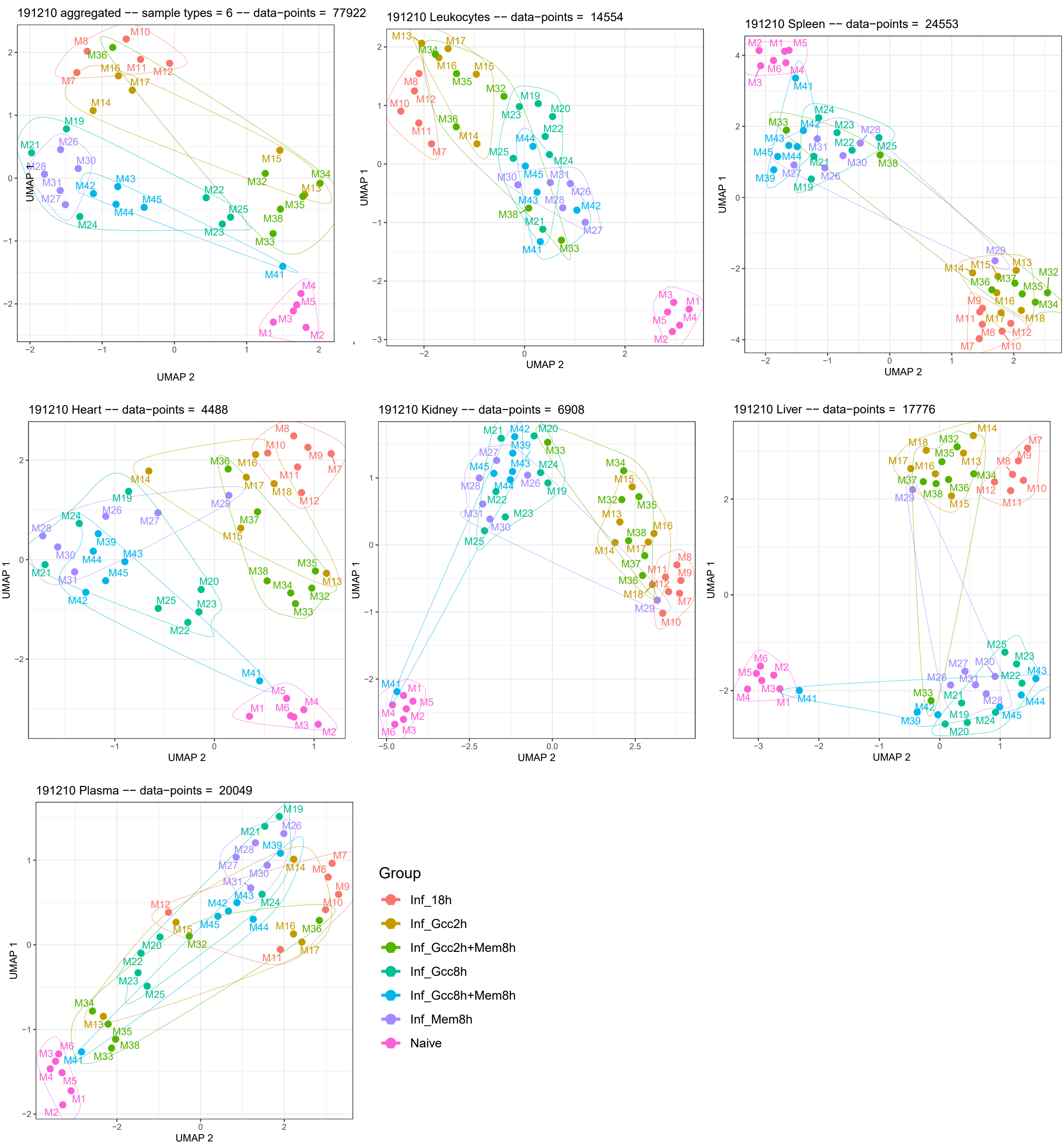


Figure S6d Highlighting intervention group clusters using UMAPs.
UMAP visualization was applied to the proteome data from all interventions across all organs and colors indicate tretament groups. Smaller clusters indicate similarity of individuals within an intervention group. Higher spread inicates dissimilar proteome states within an intervention group.
